# Adaptive cell invasion maintains organ homeostasis

**DOI:** 10.1101/2020.12.01.404954

**Authors:** Julia Peloggia, Daniela Münch, Paloma Meneses-Giles, Andrés Romero-Carvajal, Melainia McClain, Y. Albert Pan, Tatjana Piotrowski

**Affiliations:** Stowers Institute for Medical Research, Kansas City, MO 64110, USA.; Pontificia Universidad Católica del Ecuador, Escuela de Ciencias Biológicas, Quito, Ecuador.; Center for Neurobiology Research, Fralin Biomedical Research Institute at Virginia Tech Carilion, Virginia Tech, Roanoke, Virginia 24016.; Department of Biomedical Sciences and Pathobiology, Virginia-Maryland Regional College of Veterinary Medicine, Virginia Tech, Blacksburg, Virginia 24060.; Department of Psychiatry and Behavioral Medicine, Virginia Tech Carilion School of Medicine, Roanoke, Virginia 24016.

**Keywords:** lateral line system, zebrafish, ionocytes, developmental biology

## Abstract

Mammalian inner ear and fish lateral line sensory hair cells depend on fluid motion to transduce environmental signals and elicit a response. In mammals, actively maintained ionic homeostasis of the cochlear and vestibular fluid (endolymph) is essential for hair cell function and numerous mammalian hearing and vestibular disorders arise from disrupted endolymph ion homeostasis. Lateral line hair cells, however, are openly exposed to the aqueous environment with fluctuating ionic composition. How sensory transduction in the lateral line is maintained during environmental changes of ionic composition is not fully understood. Using lineage labeling, *in vivo* time lapse imaging and scRNA-seq, we discovered highly motile skin-derived cells that invade mature mechanosensory organs of the zebrafish lateral line and differentiate into Neuromast-associated (Nm) ionocytes. Furthermore, the invasive behavior is adaptive as it is triggered by drastic fluctuations in environmental stimuli. Our findings challenge the notion of an entirely placodally-derived lateral line and identify Nm ionocytes as regulators of mechanosensory hair cell function by modulating the ionic microenvironment. The discovery of lateral line ionocytes provides an experimentally accessible *in vivo* system to study cell invasion and migration, as well as the physiological adaptation of vertebrate organs to changing environmental conditions.

## Introduction

The vestibular and auditory part of the vertebrate inner ear, as well as the lateral line sensory organ of aquatic vertebrates harbor hair cells immersed in electrogenically-maintained fluid microenvironments essential for sensory transduction. Inner ear hair cells perceive sounds and changes in body position and lateral line hair cells detect water motion and enable the animals to orient themselves, catch prey and avoid predators (1). Changes in the ionic composition of the fluid environment in the mammalian inner ear leads to hearing loss and vestibular defects (2). Vestibular and auditory systems are known as closed systems because of the specialized K+-rich fluid maintained by these organs, called the endolymph. The lateral line, on the other hand, is referred to as an open system as it is devoid of endolymph and its organs are exposed to the aqueous habitat in which the fishes and amphibians reside. However, lateral line hair cells are covered by a complex gelatinous cupula that provides an electrical and ionic microenvironment that is actively maintained and necessary for the lateral line cells to depolarize when mechanically activated (3, 4). The mechanisms regulating ionic homeostasis in these open systems are not fully understood, and presently, the cell types maintaining the proper ionic composition within the cupula remain unknown (5–10). Several mechanisms that regulate ionic and osmotic homeostasis have evolved in vertebrates. For example, cells express different ion channels for fine tuning ion concentrations in their cytoplasm. In addition, kidneys, the inner ear, mammalian lungs, as well as gills and embryonic skin of fish possess specialized cells for ion uptake and secretion, some of which are called ionocytes (11). While ion-transporting cells in the kidney help maintain body ion homeostasis related to excretion, the loss of several ion channels in the inner ear causes impairment of sensory hair cell function and hearing loss (2). Likewise, both genetic ablation of skin ionocytes in zebrafish or exposure to acidified water causes impaired hair cell function, supporting the notion that lateral line hair cells also need a buffered ionic environment for proper function (12–14). To achieve homeostasis, the concentrations of multiple ions need to be regulated. In the mammalian inner ear, the mechanisms regulating the ionic composition of the endolymph involve specialized K+, Na+ and Cl− channels and transporters in the epithelial cells neighboring the hair cells (15, 16). In the skin and gills of zebrafish, five subtypes of ionocytes are described to fine tune whole-body ionic balance: NaR (Ca2+ uptake), HR (H+ secretion/Na+ uptake/NH4+ excretion), NCC (Na+/Cl− uptake), KS (K+ secretion) and SLC26 (Cl− uptake/HCO3− secretion) (17–23). Neither in the vertebrate ear or the lateral line the cellular and molecular basis of ion concentration control are completely understood.

The zebrafish sensory lateral line has proven to be an excellent model system to study sensory organ development and regeneration in vivo (1, 24) and its accessibility allowed us to discover a cell type that modulates ion concentrations and hair cell function. The superficially located mechanosensory organs, called neuromasts, contain centrally located mechanosensory hair cells that are surrounded by several types of support cells. Loss of hair cells triggers a robust regenerative response during which support cells divide and differentiate into new hair cells (25). The lateral line system, including the sensory organs, the sensory ganglia, and the axons that innervate the hair cells is derived from sets of embryonic ectodermal placodes (26, 27). In the trunk, the posterior lateral line placode (primI) migrates toward the tail tip between 24-48 hours post fertilization (hpf) and periodically deposits clusters of cells that differentiate into neuromasts. Starting at 48 hpf, a second placode (primII) migrates along the trunk and deposits neuromasts of a different orientation (28). Despite different deposition times, neuromasts from all placodes have mature and functional hair cells by 72 hpf (29, 30).

Using a combination of live imaging and single-cell RNA sequencing (scRNA-seq), we report the discovery of an invasive cell type in the zebrafish lateral line possessing ionocyte characteristics, named Nm ionocytes. Cre-mediated lineage tracing and time-lapse imaging demonstrate that Nm ionocytes are derived from *krtt1c19e*+ basal keratinocytes (31) that normally surround neuromasts. We find that these cells both migrate extensively and persistently invade the post-embryonic and adult lateral line organs. It remains unclear how prevalent non-pathological cell invasion may be in post-natal and adult tissues and whether or not such events are active contributors to normal organ function. In the lateral line the invasion process is triggered and modulated in response to changing environmental stimuli, such as salinity and pH, a process we refer to as adaptive cell invasion (ACI). The discovery of ACI and Nm ionocytes indicate important and underappreciated roles for post-embryonic developmental processes and environmental stimuli in regulating mechanosensory organs in vertebrates, including humans.

## Results

### Zebrafish neuromasts contain previously uncharacterized cells that express ionocyte markers

To characterize lateral line cell type heterogeneity in mature 5 days post fertilization (dpf) neuromasts, we previously performed scRNA-seq analysis with cells isolated via fluorescence activated cell sorting (FACS) from a double transgenic line that labels hair and support cells (Figure 1A and 1B; *Et(Gw57a:EGFP);Tg(pou4f3:GFP)*) (32). Using confocal imaging we discovered cells in some neuromasts that were not labeled by lateral line markers (Figure 1C and 1D, Supplementary Figure 1A’, 1B’, 1C’, asterisks). These cells present in these fluorescent “gaps” (usually one or two per neuromast) were therefore not sorted and included in previously published datasets. The majority of gaps contains a pair of cells, rarely three, as revealed by the ubiquitous, nuclear marker H2B-mCherry (Figure 1C-E and Supplementary Figure 1A” and B”, white dots). To identify and characterize these cells, we looked for transgenic lines that would label them. We found that the Notch pathway reporters *Tg(tp1bglobin:mCherry or EGFP)* (33), label only one of the two cells in the pair (Figure 1F and 1G, white arrowheads, Figure 1H-J, Supplementary Figure 1C”, white dot). These lines, from now on called Notch reporters, are mosaic and only label 80% of the observed gaps in fluorescence (Figure 1J, Supplementary Figure 1C, yellow arrowhead). Additionally, quantification of the lateral line specific *she* enhancer driving mCherry *(Tg(she:H2A-mCherry)*) in Notch reporter-positive (Notch+) cells confirmed that these cells are indeed not labeled by lateral line markers (Supplementary Figure 1D-F).

**Fig. 1.**
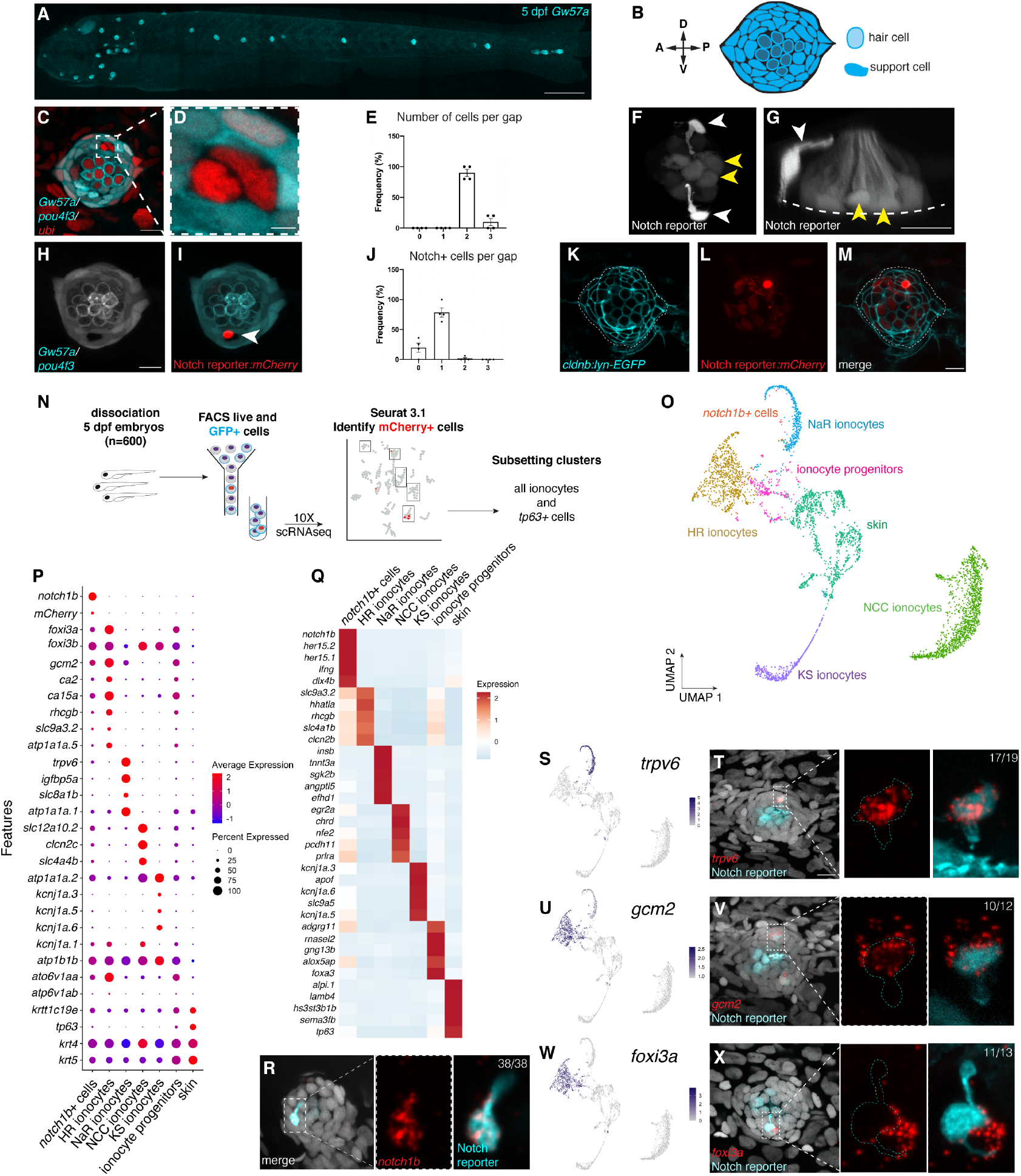
Zebrafish neuromasts contain a small subset of previously uncharacterized pairs of cells that express ionocyte markers. **(A)** Maximum projection of a confocal z-stack of a 5 days post fertilization (dpf) *Et(Gw57a:EGFP; Tg(pou4f3:GFP)* larva depicting lateral line neuromasts in cyan. Scale bar equals 300 μm. (B) Schematic of neuromast nuclei, dorsal view. **(C)** Triple transgenic larvae of *ubi:H2A-mCherry* that ubiquitously labels nuclei with the hair and support cell labeling (*Et(Gw57a:EGFP)*) and the hair cell marker *Tg(pou4f3:GFP)* shows a pair of cells that does not express neuromast or hair cell markers. Dorsal view. **(D)** Magnification of the unlabeled cell pair in (C). **(E)** Quantification of (C) showing how many cells are present per unlabeled gap (n = 4 fish). Error bars indicate standard error of the mean (SEM). **(F)** One cell of the pair is labeled by the Notch reporter line *tp1bglobin:EGFP* (white arrowheads). The Notch reporter also labels central support cells in the neuromast (yellow arrowheads). **(G)** 3D projection of a confocal z-stack of a neuromast showing one of the cells with an apical projection (white arrowhead) and central support cells (yellow arrowheads) from a lateral view (Supplementary movie 1). **(H)** Z slice of a neuromast labeled by *Et(Gw57a:EGFP);Tg(pou4f3:GFP)* and **(I)** *tp1bglobin:hmgb1:mCherry* confirms that the Notch reporter+ (Notch+) cell is one of the cells in the gap observed in (C). **(J)** Quantification of Notch+ cells per gap (n = 4 fish, 16 gaps). Error bars indicate SEM. (K-M) Double transgenic larvae *cldnb:lyn-GFP;tp1bglobin:mCherry* in which all neuromast cells and skin cells **(K)** and the Notch+ cell **(L)** and its partner cell in the pair are labeled, were used for cell sorting. **(M)** Merged images from (K) and (L). **(N)** Single-cell RNAseq (scRNA-seq) sorting strategy and flow. **(O)** UMAP of the different clusters after subsetting ionocytes and *tp63*+ cells. **(P)** Dot plot of known ionocyte markers in major ionocyte clusters and clustered *notch1b*+ cells. **(Q)** Pseudobulk heatmap of cluster markers. **(R)** *notch1b* hybridization chain reaction (HCR) confirms that notch1b is expressed in the Notch+ cell in the neuromast. **(S)** Feature plot for *trpv6*. **(T)** *trpv6* expression in the Notch+ cell in the neuromast, as shown by HCR (*n* = 6 fish, 12 neuromasts). n-numbers on the top right indicate the number of Nm ionocyte pairs analyzed. A higher magnification image of the boxed area is shown on the right. **(U)** Feature plot for *gcm2*. **(V)** *gcm2* is expressed in both cells in the pair (*n* = 4 fish, 11 neuromasts). **(W)** Feature plot for *foxi3a*. **(X)** *foxi3a* is expressed in the Notch− ionocyte (*n* = 5 fish, 12 neuromasts). All images except panel A and G were blurred. Panel A, F and G had non-linear adjust (gamma) to allow for visualization of the skin of the larvae and all labeled cells in the neuromast, respectively. Scale bars equals 10 μm unless otherwise noted.

To determine the identity of the pairs of non-lateral line cells, we performed scRNA-seq of cell purified with FACS from double transgenic *Tg(cldnb:lyn-GFP)*;Notch reporter:mCherry fish (Figure 1K-M). The *cldnb* promoter labels lateral line neuromasts, skin cells, as well as the Notch+ and Notch− non-lateral line cells, capturing both cells when sorted by GFP expression. In contrast, GFP and mCherry are co-expressed only in Notch+ non-lateral line cells, in neuromast central support cells and rarely in some skin cells (Supplementary Figure 1G and 1G’). To capture non-lateral line cells within the mature neuromasts, we selected 5 dpf fish, in which around 50% of neuromasts have fluorescent gaps with pairs of these non-lateral line cells. This would also allow us to directly compare our results with previous published neuromast datasets and use mCherry detection in central support cells as a positive control (32). We performed mild larvae dissociation to avoid contamination from deeper tissues and sorted for live GFP+ cells (Figure 1N). After sequencing, alignment and quality controls, we performed dimensionality reduction with UMAP (Supplementary Figure 1H, Supplementary Table 1) (34). To identify the Notch+ non-lateral line cells we searched the dataset for the presence of mCherry mRNA-expressing cells. We identified most mCherry transcripts in clusters expressing markers for NCC, NaR and HR ionocytes and, as expected, in the neuromast clusters containing central support cells (Supplementary Figure 1I).

To gain a better insight on the identity of mCherry+ cells, we selected for further analysis all detected ionocyte clusters together with *tp63*+ skin stem cells that contain ionocyte progenitors during embryonic development (31, 35, 36). This generated clusters of four major known ionocyte sub-types, skin cells and putative ionocyte progenitor populations (Figure 1O). We annotated ionocyte subtype clusters based on their expression of classical markers: *trpv6*, *igfbp5a* and *slc8a1b* are expressed in NaR ionocytes, *rhcgb*, *slc4a1b*, *foxi3a*, *ca2* and *atp6v1aa* in HR ionocytes, *clcn2c*, *slc4a4b* and *slc12a10*.2 in NCC ionocytes and *kcnj1a* channels in KS ionocytes (18, 37). *krtt1c19e* and *tp63* expressing clusters were combined and identified as “skin” cells, while clusters that form streams between the skin and HR ionocytes were annotated as putative ionocyte progenitors, because of their expression of basal keratinocyte (*tp63*, *krtt1c19e*, *krt4*), ionocyte progenitor (*foxi3a/b*) and ionocyte markers (*ca15a*, *atp1b1b*, *atp6v1aa*) (18, 31, 35, 36). The mCherry+ cells did not naturally fall into a homogenous cluster. To be able to perform differential expression analyses between Notch reporter cells and ionocytes, we selected and clustered all mCherry+ cells that were originally intermingled with ionocyte cells in UMAP space. The most highly expressed Notch receptor in these cells was *notch1b* (Figure 1P and 1Q, “*notch1b*+ cells”). To test if the non-lateral line Notch+ cells express *notch1b*, we performed hybridization chain reaction (HCR) with *notch1b* in Notch reporter:EGFP larvae (38). Indeed, *notch1b* mRNA colocalized with EGFP, suggesting that cells in the *notch1b* cluster possibly represent one of the cells in the pair of non-lateral line cells in the neuromast (Figure 1R). In agreement with this finding, Notch+ non-lateral line cells in neuromasts were detected by anti-Notch1 immunofluorescence (Supplementary Figure 1J).

Strikingly, our scRNA-seq analysis showed that some traditional ionocyte markers are not as specific for ionocytes subtypes as previously assumed (Figure 1P). We now identified highly specific marker genes for these cell types (Figure 1Q and Supplementary Table 2) (18). Likewise, *notch1b*+ cells express specific genes that they do not share with ionocytes, such as *her15.1*, *her 15.2* and *lfng* (Figure 1Q). However, they also express a few NaR and HR ionocyte-associated genes, albeit at a reduced level. Examples are the calcium channel *trpv6* expressed in NaR cells and the transcription factor *gcm2* that has been associated with both NaR and HR ionocyte specification (Figure 1P, 1S-V) (39, 40). On the other hand, the Notch+ cell does not express sodium-potassium channels based on immunohistochemistry of Na+/K+-ATPases (NKA), labeling NaR and HR cells (Supplementary Figure 1K, K’).

We could not distinguish the non-lateral line Notch− cells from skin ionocytes in our scRNA-seq data. Possibly, they failed to form a distinct cluster because we sequenced too few cells, or their transcriptional signature is too similar to skin ionocytes. HCR with candidate genes revealed that the Notch− non-lateral line cells, as well as skin ionocytes strongly express the HR/NaR specification gene *gcm2* and the HR ionocyte marker *foxi3a* (Figure 1U-X, Supplementary Figure P). Additionally, the transcription factor *foxi3b* and the NaR marker *atp1a1a.1* are likely expressed in either of these cells (Supplementary Figure 1L-O). Neither the Notch+ nor the Notch− non-lateral line cells express markers for NCC or KS ionocytes (Supplementary Figure 1P, Q). We therefore conclude that the Notch− cell in the pair of unlabeled cells we discovered in neuromasts is an ionocyte that expresses both HR and NaR markers. The Notch+ cell expresses a few ionocyte-associated genes but otherwise does not share the majority of ionocyte marker genes. We therefore hypothesize that the Notch+ cell is likely an accessory cell as has been described for gill ionocytes (41–44). We termed the pair of cells Neuromast-associated ionocytes (Nm ionocytes).

### Nm ionocytes are derived from skin cells surrounding the neuromast

As Nm ionocytes share genes with skin ionocytes but do not express neuromast markers we wondered if they possess a different embryonic origin than the placodally-derived neuromast cells (Figure 2A). To determine the embryonic origin of these cells, we analyzed transgenic zebrabow fish that allow tracing of cells throughout development (45). Briefly, after Cre-induced recombination of a set of mutually incompatible loxP sites, only one out of three fluorescent proteins per construct (RFP, CFP or YFP) is stochastically expressed in each cell. Fish lines containing more than one insertion produce a combination of colors with clonally related cells having the same color (Figure 2B). Cre-induced recombination between 4-6 hpf led to mosaically labeled lateral line primordia which then deposited mosaic neuromasts (Figure 2C-E). When we followed these neuromasts over time we detected two changes: 1) Starting between 48-72 hpf, some neuromasts suddenly contained pairs of cells in the poles that had very different colors than the other cells in the neuromast (Figure 2E) and 2) The vast majority of neuromast cells became clonal in adult fish, highlighting the presence of differently labeled pairs (Figure 2F, F’, arrowheads). Triple transgenic embryos for *Tg(ubi:Zebrabow;ubi:creERt2)* and a Notch reporter driving the expression of mKate2 revealed that one of the differently colored cells in each pair is mKate2+, demonstrating that the differently colored pairs of cells are Nm ionocytes (Supplementary Figure 2A). Thus, Nm ionocytes possess a different embryonic origin than all other placodally-derived neuromast cells.

**Fig. 2.**
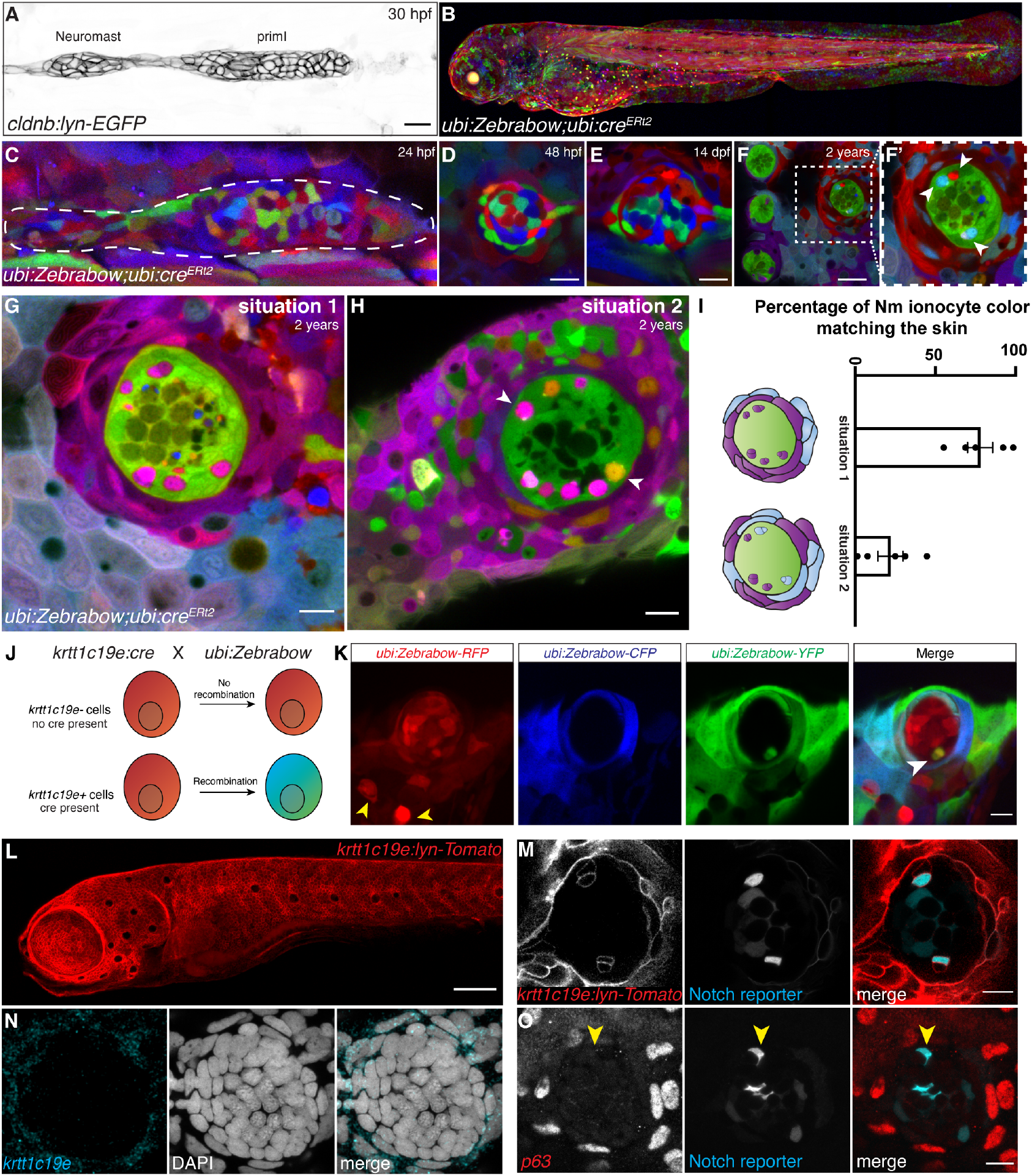
Neuromast-associated ionocytes are derived from skin cells immediately surrounding the neuromast. **(A)** Lateral line primordium depositing a neuromast at 30 hours post-fertilization labeled by the transgenic line *cldnb:lyn-GFP*. **(B)** Maximum intensity projection of a 3 dpf *ubi:Zebrabow* larvae recombined with 4-OHT during shield stage. **(C)** Lateral line primordium of a 24 hpf fish recombined with 4-OHT (15 min at 4 hpf) shows a mosaically labeled primordium. Same neuromast deposited from primordium in (C) is still mosaic at 48 hpf **(D)** and 14 dpf **(E)**. **(F)** Neuromasts in two-year-old fish that were recombined between 16- 24 hpf are no longer mosaic and have become clonal. **(G)** Neuromasts of recombined *ubi:Zebrabow;ubi:creERT2* larvae possess Nm ionocyte pairs of the same color hue as some skin cells surrounding the neuromast. All other neuromast cells have the same color. **(H)** Neuromasts with Nm ionocyte pairs of two different color hues (white arrows), but still sharing color hues with surrounding skin cells. **(I)** Quantification of the percentage of neuromasts in which all Nm ionocyte pairs possess the same color, situation 1, or the percentage of neuromasts in which the pairs are not all of the same color, situation 2 (n = 5 fish, 194 neuromasts). Error bars indicate SEM. **(J)** Overview of lineage tracing experiments with *ubi:Zebrabow* and *krtt1c19e:cre-MYC*. *krtt1c19e*+ basal keratinocytes express constitutively active Cre recombinase enzyme, which recombines *ubi:Zebrabow* in basal keratinocytes and their progeny, changing their color from red to CFP or YFP. **(K)** Neuromast of 5 dpf *ubi:Zebrabow;krtt1c19e:cre-MYC* larva shows a pair of recombined cells (white arrowhead) in between red, unrecombined neuromast cells. Skin ionocytes differentiate before Cre expression and are not recombined (yellow arrowheads). **(L)** Overview of 5 dpf larvae from transgenic line *krtt1c19e:lyn-Tomato*, that labels basal keratinocytes. **(M)** Neuromast of 5 dpf *krtt1c19e:lyn-Tomato;tp1bglobin:EGFP* fish shows pairs of *krtt1c19e*+ cells inside the neuromast, one of which is also labeled by the Notch reporter. *krtt1c19e* channel had gamma adjusted to allow for observation of weakly labeled cell types. **(N)** *krtt1c19e* HCR and **(O)** TP63 immunohistochemistry do not label cells in the neuromast, suggesting Nm ionocytes no longer express the mRNA of skin markers. Scale bars in all images equal 10 μm.

We also observed that Nm ionocytes shared their color hue with skin cells immediately surrounding the neuromasts (Figure 2G and 2H), suggesting that skin cells might be the source of the Nm ionocytes, or at least share the same origin. In the majority of neuromasts (84%), all ionocytes within a neuromast shared the same color hue, while in 16% of neuromasts ionocytes had two or more different color hues (Figure 2I). Irrespective of Nm ionocyte color heterogeneity, we always observed that cells in the surrounding skin expressed the same color hues. We did not observe any neuromasts in which the ionocyte color was not shared by surrounding skin cells (*n* = 194 neuromasts). As Nm ionocytes often exist in pairs of cells we wondered if the two cells are clonally related. While 87% of pairs had cells with the same color, 13% of pairs were composed of cells of different colors (Supplementary Figure 2B and 2C). However, in the case of the differently colored pairs we cannot exclude that they were derived from a common progenitor cell but that the recombination event occurred after division of this progenitor cell generating differently colored sister cells.

To investigate if Nm ionocytes are derived from skin cells, we performed lineage tracing by crossing the *ubi:Zebrabow* fish to *Tg(krtt1c19e:cre-MYC)*, that expresses a constitutively active Cre recombinase with a MYC-tag (Figure 2J). The *krtt1c19e* promoter becomes active around 24 hpf, permanently labelling basal keratinocytes and their progeny. Skin ionocytes already differentiate around the 14-somite stage (16 hpf) and are therefore not recombined (Figure 2K, yellow arrows). We observed recombined pairs of cells in the neuromasts, suggesting they are derived from Cre-expressing basal keratinocytes (Figure 2K, white arrow). Likewise, double transgenic larvae for the basal keratinocyte reporter line *Tg(krtt1c19e:lyn-Tomato)* (Figure 2L) and the Notch reporter expressing EGFP revealed pairs of Tomato+ cells in the neuromasts, of which one cell expressed the Notch reporter, confirming that Nm ionocytes initially expressed the basal keratinocyte cell marker *krtt1c19e* (Figure 2L and 2M). Yet, *krtt1c19e* mRNA and TP63 protein are absent from Nm ionocytes suggesting these markers are downregulated as they differentiate (Figure 2N and 2O). In support of this hypothesis, we found a negative correlation between fluorescence of *krtt1c19e* and Notch reporter expression (Supplementary Figure 2D and 2E), suggesting that the cells stop expressing *krtt1c19e* as Notch signaling increases. Altogether, we conclude from these data that Nm ionocytes are likely derived from basal keratinocytes surrounding the neuromasts and differentiate once inside the sensory organ.

### Nm ionocyte progenitors migrate into neuromasts as pairs of cells

Nm ionocytes are not present at the time of neuromast deposition. To investigate the origin and dynamics of Nm ionocytes *in vivo* we took advantage of the superficial location of the lateral line. Since we detected the first Nm ionocytes around 3 dpf, we performed time lapse imaging starting at this developmental stage to capture the moment when these cells first appeared. Time lapse recordings revealed that ionocyte progenitors migrate into neuromasts as pairs of cells (Figure 3A and B, colored dots, Supplementary Movies 2 and 3). By tracking individual pairs of Nm ionocytes from their origin outside the neuromast (Figure 3C) to their final location (Figure 3D, Supplementary Movie 4), we found that after invasion some pairs of cells migrate to the other side of the organ (Figure 3C-E, Track 1 and 2), while others stay closer to their site of entry (Figure 3C-E, Track 3 and 4). Quantifying the speed of Notch+ cells revealed that these cells on average migrate more than seven times the distance from their start to final location (Figure 3F). Independently from the distance migrated, both cells re-arrange extensively between neuromast support cells and hair cells, while remaining closely associated with each other.

**Fig. 3.**
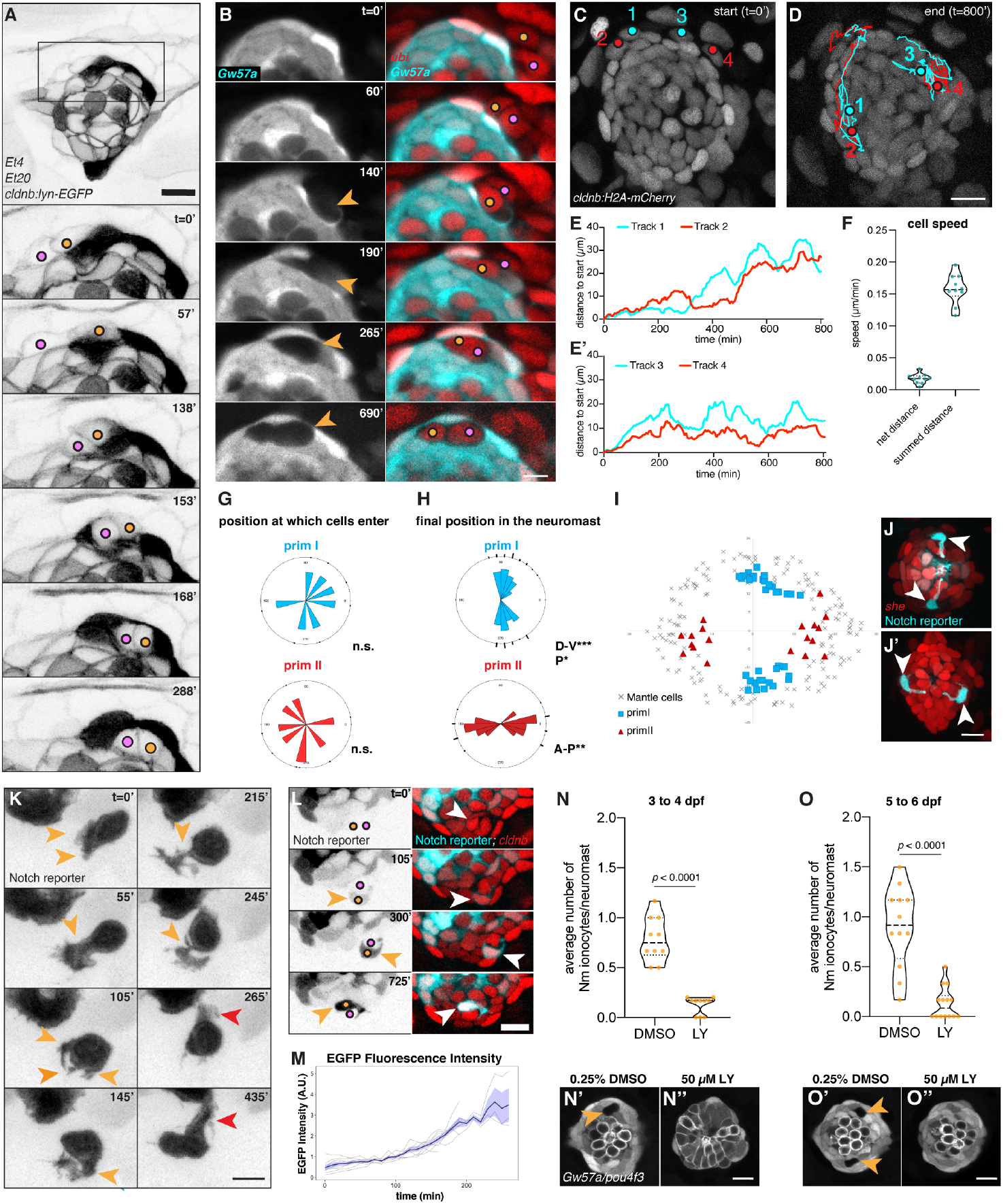
Ionocytes move into the neuromast as rearranging pairs of cells and their migration depends on Notch signaling. (A) Still images of a time lapse recording depicting two Nm ionocyte precursors (colored dots) entering a neuromast of a 3 dpf *Et4;Et20;cldnb:lyn-EGFP* larva (Supplementary Movie 2). Top image shows an overview of the neuromast at the start of the time lapse with boxed area labeling the region shown at higher magnification below (scale bar = 5 μm). Images were gamma-adjusted to allow for better visualization of the cell membranes. **(B)** Individual z-slices of a time lapse recording of a *Et(Gw57a:EGFP);ubi:H2A-mCherry* larva at 3 dpf, showing Nm ionocyte precursor cells migrating into the neuromast as visualized by translocation of the two cell nuclei (right panels, colored dots) as well as the appearance of a gap in *Et(Gw57a:EGFP)* fluorescence (left panels, orange arrowheads, Supplementary movie 3). Scale bar = 5 μm. **(C-D)** Example tracks of two Nm ionocyte precursor cell pairs (cell tracks 1-4) in a 2 dpf *cldnb:H2A-mCherry* larva entering the neuromast and re-arranging extensively (Supplementary movie 4). Red and cyan dots depict the position of the Notch− as well as the future Notch+ cell of each pair at the start (C) and end point of the recording (D), respectively. **(E-E’)** Quantification of the distance from the start position of the cells tracked in C and D over time shows both cells of each pair migrating and re-arranging synchronously, in close association to each other. **(F)** Quantification of the cell speed calculated based on the net distance, as well as the summed distance that Notch+ cells traverse, respectively (*n* = 7 neuromasts, 10 cells). Rose plots displaying the angular position at which Nm ionocyte precursors enter neuromasts of different axial polarities (primI- and primII-derived neuromasts with blue and red labeling, respectively) between 2 and 3 dpf (*n* = 15 neuromasts, 7 fish). Binomial test revealed no directional bias. **(H-I)** Angular, final position of Nm ionocytes in primI- and primII-derived 5 dpf neuromasts (H), and their spatial distribution in relation to mantle cell position (I, *n* = 55 neuromasts). Binomial analysis revealed significant bipolar localization with a slight directional bias towards posterior in primI-derived neuromasts. **(J-J’)** Representative images of Nm ionocyte morphology and localization in primI-derived (J) and primII-derived neuromasts at 5 dpf (J’). Images were gamma-adjusted to allow better visualization of fine cellular projections. **(K)** Still images of a time lapse recording at 3 dpf, depicting a Notch+ Nm ionocyte precursor cell extending multiple short-lived protrusions (orange arrowheads) before stabilizing a major extension apically (red arrowheads, Supplementary Movie 5). Images were gamma-adjusted to allow better visualization of fine cellular projections. Scale bar = 5 μm. **(L)** Gradual increase in Notch reporter fluorescence (left panel, orange arrowheads; right panel, white arrowheads) as both Nm ionocyte precursor cells (left panel, colored dots) re-arrange between hair cells and support cells (Supplementary movie 6). **(M)** Quantification of Notch reporter GFP fluorescence over time, normalized by the average of all the curves up to the 160 min time point. **(N-N”)** Inhibition of Notch signaling with 50 μM gamma secretase inhibitor LY411575 (*n* = 10 larvae, 60 neuromasts) for 24h between 3 and 4 dpf increases hair cell numbers at the expense of support cells and drastically reduces Nm ionocyte frequency based on gaps in *Et(Gw57a:EGFP)* fluorescence compared to DMSO controls (*n* = 10 larvae, 60 neuromasts; Mann-Whitney test, p < 0.0001). Representative images shown below (N’ and N”). **(O-O”)** Inhibition of Notch signaling with 50 μM LY411575 for 24h (*n* = 14 larvae, 84 neuromasts) between 5 and 6 dpf still abolishes Nm ionocyte frequency based on Et(Gw57a:EGFP) fluorescence compared to DMSO controls (*n* = 12 larvae, 72 neuromasts; Mann-Whitney test, p < 0.0001). Representative images shown below (O’ and O”). Scale bars of all images = 10 μm unless otherwise noted. Dashed black lines in violin plots indicate the median, while dotted black lines indicate quartiles.

Once inside the neuromast, the Notch+ ionocyte precursor extends highly dynamic protrusions as visualized by the cytoplasmic Notch reporter (Figure 3K, orange arrowheads, Supplementary Movie 5). While the time lapse recording showed that Nm ionocyte precursors do not enter the organs at predefined positions (Figure 3G), spatial analyses of Nm ionocytes at 5 dpf demonstrate that these cells are eventually positioned stereotypically in the D-V and A-P poles of primI- and primII-derived neuromasts, respectively (Figure 3H and 3I, blue squares and red triangles, respectively, Figure 3J and 3J’, arrowheads). Once the pair reaches the poles, Nm ionocytes stop re-arranging and project a stable apical extension towards the cuticular plate of the neuromast (Figure 3K, red arrowheads).

Notch signaling plays a crucial role in skin ionocyte specification via lateral inhibition [30] and we observed that migration and differentiation of Nm ionocytes are associated with a progressive increase in Notch signaling in one cell of the pair (Figure 3L, arrowheads and Figure 3M; Supplementary Movies 6 and 7). We therefore wondered if inhibiting Notch signaling would affect the fate of the cells in the pair. As Notch inhibition abrogates the expression of the fluorescent Notch reporter, we analyzed neuromasts in which support and hair cells were labeled and Nm ionocytes appear as “gaps” in between fluorescently labelled cells (Figure 3N’ and 3N’, arrowheads; *Et(Gw57a:EGFP);Tg(pou4f3:GFP)*). Pharmacological inhibition of Notch signaling with the gamma secretase inhibitor LY411575 during neuromast development from 3 to 4 dpf generated an excess of hair cells at the expense of support cells accompanied by a lack of ionocytes (Figure 3N, 3N’ and 3N”) [7]. To exclude the possibility that the excess of hair cells causes crowding and spatially prevents ionocyte precursors from invading the neuromast, we next inhibited Notch signaling once all hair cells had differentiated (5 dpf) and some Nm ionocytes are already present in the neuromast. We still observed a significant decrease in Nm ionocytes (Figure 3O, 3O’ and 3O”). We conclude from these data that Notch signaling likely plays an essential role not only in the migration and differentiation of the highly protrusive ionocyte progenitors invading mature neuromasts as pairs, but also in their survival as Notch inhibition between 5 and 6 dpf abolishes pre-existing Nm ionocytes.

### 3D modeling of Nm ionocytes reveals exquisite morphological characteristics resembling gill and skin ionocytes

Nm ionocyte differentiation is associated with profound changes in cell morphology. Although these cells commonly exist as pairs, we could only visualize the Notch+ cell during live imaging experiments due to a dearth of markers. To visualize the structure of both cells, we performed Serial Block Face Scanning Electron Microscopy (SBF-SEM) to 3D reconstruct both cells. To identify the ionocyte pair inside the neuromast, we took a correlative approach between SBF-SEM and light microscopy. We imaged ionocyte-containing neuromasts under a confocal microscope with two fluorescent markers: a neuromast nuclei marker (red) and the Notch reporter (cyan) (Figure 4A). The same embryo was then processed for SBF-SEM. Modeling of the SEM neuromast nuclei resulted in a very similar pattern of nuclei observed under confocal microscopy (Figure 4B) allowing us to identify the nuclei of the ionocyte pairs in the SBF-SEM dataset (Figure 4A and 4B). 3D modeling of the cell membranes of the Notch+ and Notch− cells revealed the tight association between these cells and their morphology. The two cells are in close contact with each other along their entire apico-basal axis, their nuclei are located basally and the Notch+ cell (cyan) is smaller than the Notch− cell (white) (Figure 4C, Supplementary Movie 8). Both cells possess extensions that reach the apical surface of the neuromast and frequently contain thin projections (Figure 4C and 4D, Supplementary Figure 3A and 3A’, arrowheads).

**Fig. 4.**
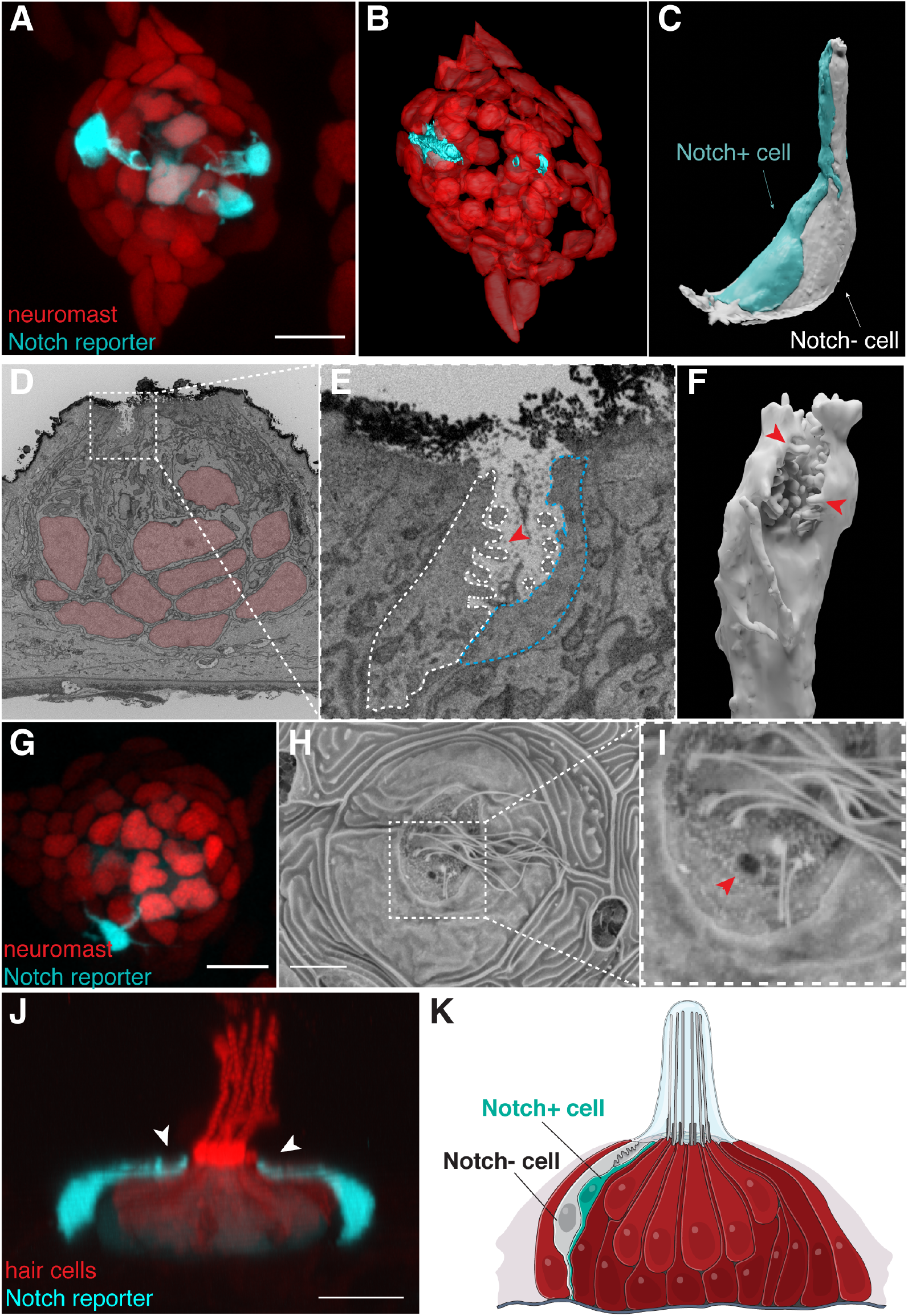
Neuromast-associated ionocytes share morphological characteristics with skin ionocytes and are exposed to the external environment through an opening to the neuromast cupula. **(A)** Maximum intensity projection of a neuromast labeled by *she:H2A-mCherry;tp1bglobin:EGFP* shows nuclei of the neuromast cells in red and three Notch+ Nm ionocytes in cyan. **(B)** 3D modeling of the SBF-SEM stack of the same neuromast shown in (A). **(C)** 3D modeling of the Nm ionocyte pair shows intricate connections between the two cells, with a pronounced apical extension. **(D)** Single image of a transverse section of the SBF-SEM shows the apical part of a pair of Nm ionocytes (white square) and neuromast cells (red nuclei). **(E)** Magnification of white square in (D) showing microvilli (red arrowhead) and the apical crypt of the ionocyte pair exposed to the outside of the neuromast. **(F)** 3D modeling of the microvilli (red arrowheads) containing crypt of the Notch− cell. **(G)** Maximum intensity projection of a neuromast labeled by *she:H2A-mCherry;tp1bglobin:EGFP.* **(H)** Scanning electron micrograph of the same neuromast shown in (G). **(I)** Magnification of the region in white square in (H). Red arrowhead depicts an opening in the neuromast cuticular plate that correlates with the position of the Nm ionocyte pair. **(J)** Lateral view of a confocal z-stack 3D projection of a *tp1bglobin:EGFP;Myo6b:Lck-mScarlet-I* neuromast (Supplementary movie 9) . Nm ionocytes have long apical projections with the apical crypt (white arrowheads) close to the cuticular plate. **(K)** Model of the neuromast (red) containing a pair of Nm ionocytes (white and cyan).

The pair forms a crypt at the top of the apical extensions with microvilli projecting only from the Notch− cell into the lumen (Figure 4E and 4F, Supplementary Figure 3B and 3B’). This morphology is reminiscent of gill and skin ionocytes that possess apical crypts exposed to the outside [36, 37]. To test if Nm ionocytes also open to the external environment, we analyzed correlated SEM and confocal images. Indeed, we observed openings in the neuromast cuticular plate that correlated with presence and location of Nm ionocytes (Figures 4G-I, red arrowhead). Consequently, the crypts are exposed to the gelatinous cupula microenvironment that covers the cuticular plate and hair cell cilia of live animals (Figure 4J, Supplementary Movie 9).

Microvilli-containing ionocytes accompanied by accessory cells have been described mostly in seawater fish (42). We hypothesize that the Notch+ cells act as accessory cells for the Notch− Nm ionocyte. While the function of accessory cells is not fully understood, they have been proposed to modulate paracellular pathways for ionic excretion through leaky junctions with ionocytes (46). Our results show that the pair of Nm ionocyte precursors differentiates into a microvilli-containing ionocyte and a slightly smaller accessory cell that together form a crypt open to the neuromast cuticular plate. This finding reveals a previously unappreciated cell type diversity in the neuromast (Figure 4K).

### Nm ionocyte frequency is modulated by changes in salinity and pH

The finding that apical openings of Nm ionocytes are in direct contact with the overlying cupula suggests that they could be involved in controlling the ionic microenvironment surrounding hair cells. However, we observed that not all neuromasts contain ionocytes during the larval stages. To test whether Nm ionocyte frequency increases at later time points, we quantified Nm ionocyte numbers in 14 dpf larvae and 2-year-old adult fish. Indeed, the percentage of ionocyte containing neuromasts (Figure 5A), as well as the number of Nm ionocytes per neuromast (Figure 5B and 5C) significantly increases not only during early larval development (3 dpf to 5 dpf) but also from later larval stages (14 dpf) until adult stages (2 years). While we cannot fully rule out the possibility of proliferation of existing Nm ionocytes during these later stages, we observed Nm ionocytes appear in neuromasts without pre-existing ionocytes between 21 dpf and 28 dpf based on Zebrabow analysis (Figure 5D). This suggests that Nm ionocytes continue to invade neuromasts as animals age.

**Fig. 5.**
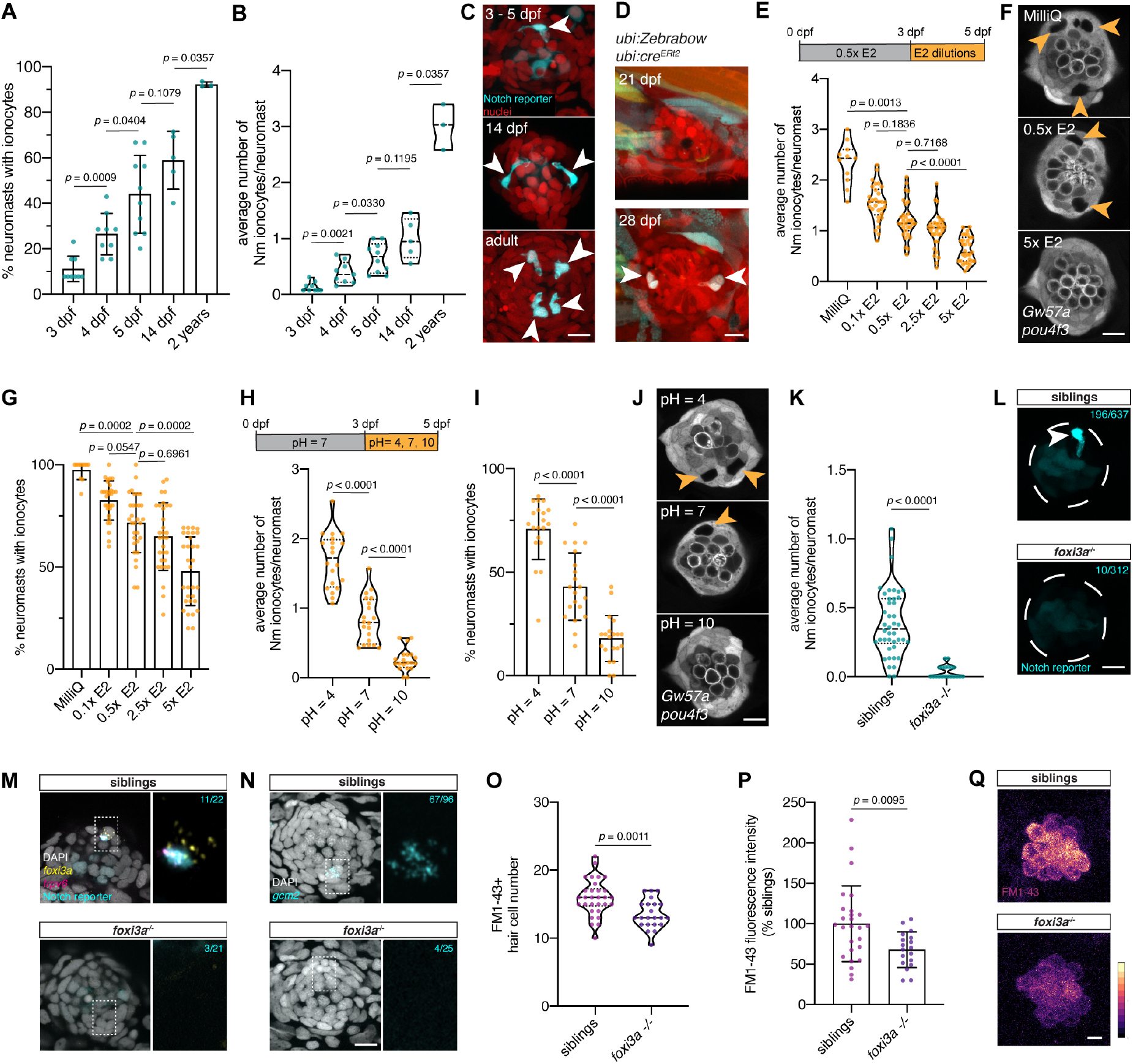
Nm ionocyte frequency is modulated by environmental changes. **(A)** The percentage of neuromasts with one or more Notch+ Nm ionocytes per fish increases between 3 dpf (*n* = 9 fish, 117 neuromasts), 4 dpf (*n* = 9 fish, 125 neuromasts), 5 dpf (*n* = 10 fish, 162 neuromasts), 14 dpf (*n* = 5 fish, 49 neuromasts) and 2 year old adult fish (*n* = 3 fish, 78 neuromasts, Mann-Whitney test). **(B)** The average number of Notch+ Nm ionocytes per neuromast increases significantly from larval up to adult stages (same n-numbers as in (A), Mann-Whitney test). **(C)** Maximum projections of confocal z-stacks depicting representative numbers of Notch+ Nm ionocytes (white arrowheads) at different stages. **(D)** Maximum projections of multiple z-slices of the same *ubi:Zebrabow;ubi:cre^ERt2^* neuromast at 21 dpf (upper panel) and 28 dpf (lower panel). Arrowheads point at newly appeared pairs of Nm ionocytes with different color hues compared to neuromast cells between 21 and 28 dpf. **(E)** Nm ionocyte number (quantified by the number of gaps in *Et(Gw57a:EGFP)* fluorescence) significantly decreases in larvae incubated in increasing salt concentrations for 48 h (MilliQ, *n* = 11 fish, 154 neuromasts; 0.1x E2, *n* = 30 fish, 433 neuromasts: 0.5x E2, *n* = 31 fish, 436 neuromasts; 2.5x E2, *n* = 31 fish, 449 neuromasts; 5x E2, *n* = 31 fish, 454 neuromasts; Kruskal-Wallis ANOVA and Dunn’s post hoc test). **(F)** Representative images of neuromasts of *Et(Gw57a:EGFP);Tg(pou4f3:GFP)* larvae that were incubated in MilliQ water, 0.5x E2 and 5x E2 medium for 48 h. Orange arrowheads point towards Nm ionocytes as visualized by gaps in the *Et(Gw57a:EGFP)* fluorescence. **(G)** The percentage of neuromasts containing one or more Nm ionocytes drastically increases in conditions of low salinity but decreases in high salinity media (same n-numbers as in (E), Kruskal-Wallis ANOVA and Dunn’s post hoc test). **(H)** Nm ionocyte frequency significantly increases in acidic conditions (pH = 4; *n* = 20 fish, 286 neuromasts) and decreases in alkaline embryo media (pH = 10; *n* = 20 fish, 291 neuromasts) compared to controls (pH = 7, *n* = 20 fish, 296 neuromasts; unpaired t-test). **(I)** The percentage of neuromasts with one or more Nm ionocytes increases with decreasing pH (same n-numbers as in (H), Mann-Whitney test). **(J)** Representative images of neuromasts of *Et(Gw57a:EGFP);Tg(pou4f3:GFP)* larvae incubated in acidic, neutral and alkaline embryo media for 48 h. Orange arrowheads indicate Nm ionocytes as visualized by gaps in the *Et(Gw57a:EGFP)* fluorescence. **(K)** The average Nm ionocyte frequency is significantly decreased in *foxi3a*^−/−^ mutants. (siblings: *n* = 42 fish, 637 neuromasts; *foxi3a*^−/−^, *n* = 20 fish, 312 neuromasts; Mann-Whitney test). **(L)** Representative images of Nm ionocytes (white arrowheads) in foxi3a mutants and siblings, as visualized by the Notch reporter. **(M)** *foxi3a* and *trpv6* HCR in *foxi3a*^−/−^ fish (lower panel, *n* = 21 neuromasts, 4 fish) and their siblings (upper panel, *n* = 22 neuromasts, 7 fish). Cyan n-numbers indicate the number of neuromasts with Nm ionocytes based on Notch reporter fluorescence as well as the expression of *trpv6*. A higher magnification image of the boxed area is shown on the right, with scale bars equaling 2 μm. **(N)** *gcm2* HCR in *foxi3a*^−/−^ fish in the Notch reporter background (lower panel, n = 25 neuromasts, 3 fish) and their siblings (upper panel, *n* = 96 neuromasts, 12 fish). Cyan n-numbers indicate the number of neuromasts with one or more Nm ionocytes based on the expression of *gcm2*. A higher magnification image of the boxed area is shown on the right. **(O)** FM1-43+ hair cell numbers are significantly reduced in *foxi3a* mutants (*n* = 15 fish, 30 neuromasts) compared to their siblings (*n* = 11 fish, 22 neuromasts). (P) The FM1-43 intensity of hair cells is significantly reduced in *foxi3a* mutants (*n* = 9 fish, 18 neuromasts) compared to their siblings (*n* = 13 fish, 26 neuromasts; unpaired t-test). Data is shown as fluorescence intensity/background, normalized to the average of the siblings and shown as percentage. **(Q)** Representative images of FM1-43 dye uptake by hair cells in *foxi3a* mutants and their siblings, as quantified in **(P)**. Scale bars equal 5 μm. All scale bars equal 10 μm, unless otherwise noted. Error bars indicate standard deviation. Dashed lines in violin plots indicate the median, dotted lines indicate quartiles. Individual data points in A, B, E, G, H, I and K each represent an average of all neuromasts quantified per fish.

The increase in Nm ionocyte frequency not only correlates with developmental age but also with the transfer of the larvae from ion-rich embryo medium (E2) to more ion-poor water in the housing facility. Since the density of skin and gill ionocytes is modulated in response to different ionic environments (47), we wondered whether salinity changes could influence the frequency of Nm ionocyte invasion. To test this hypothesis, we incubated larvae in different dilutions of E2 medium between 3 and 5 dpf and quantified the frequency of Nm ionocytes. Indeed, the number of Nm ionocytes negatively correlated with increasing E2 media salinity (Figure 5E and 5F, orange arrowheads). After incubation in MilliQ water, Nm ionocytes displayed frequencies that more closely resembled those observed in adult fish (Figure 5B and 5E). Similarly, 98.7% percent of neuromasts now contained one or more Nm ionocytes (Figure 5G). The increase in Nm ionocytes was not caused by a systemic increase in proliferation as the total neuromast cell number, as well as the number of hair cells simultaneously decreased in MilliQ water (Supplementary Figure 4A-C). Similar to Nm ionocytes, the density of skin ionocytes on the yolk increased in MilliQ water, as detected by NKA immunohistochemistry, as well as by *in situ* hybridization with the HR and NaR markers ca2 and trpv6, respectively (Supplementary Figure 4D-G). These findings provide additional support for the hypothesis that Nm ionocytes respond to salinity changes and are involved in regulating ion homeostasis.

Since incubation in MilliQ water increased Nm ionocytes but also decreased hair cell numbers, we aimed to test if hair cell death modulates Nm ionocyte numbers. Killing hair cells with the antibiotic neomycin and subsequent regeneration had no effect on Nm ionocyte frequency (Supplementary Figure 4H and 4I). Neomycin-induced hair cell death also did not lead to proliferation of the Notch+ Nm ionocyte as revealed by EdU-incorporation experiments (Supplementary Figure 4J and 4K). These results demonstrate that invasion of the neuromast by ionocytes is not triggered by hair cell death. Lateral line hair cell function is impaired in *merovingian* mutants that harbor a mutation in *gcm2* (14), a transcription factor important for the production of HR and possibly a subset of NaR ionocytes (39, 40). This impairment had been attributed to an acidification of the extracellular environment of hair cells caused by loss of skin HR ionocytes, as Nm ionocytes had not been discovered (14). Given the physical proximity to hair cells and the expression of HR ionocyte markers like *gcm2* (Figure 1V), we wondered whether Nm ionocytes could be involved in maintaining the pH in the hair cell microenvironment. We reasoned that Nm ionocyte numbers would increase in acidified environments, as it has been described for HR cells (48). Indeed, we observed that Nm ionocyte number as well as the percentage of neuromasts with ionocytes increased with decreasing pH (Figure 5H-J). This finding suggests that Nm ionocytes invade neuromasts not only in response to changes in salinity, but also pH, pointing towards an additional involvement in acid/base secretion (48).

### Loss of *foxi3a* reduces Nm ionocyte numbers and impairs hair cell function

To functionally test Nm ionocyte function we sought to identify genes that when knocked out cause Nm ionocyte loss. One candidate is *foxi3a* that, among others, is a transcriptional master regulator that specifies skin ionocytes from epidermal precursors. Morpholino-mediated knockdown results in a complete loss of NaR and HR ionocytes (35, 36). To determine whether Nm ionocytes are specified by the same transcriptional program, we generated a zebrafish mutant lacking the *foxi3a* promoter region, as well as part of the first exon by CRISPR/Cas12a-mediated genome editing (49) (Supplementary Figure 4L). Homozygous *foxi3a*^−/−^ larvae showed a significant reduction of Nm ionocytes based on Notch reporter fluorescence compared to their siblings (Figure 5K and 5L). Likewise, HCR revealed a strong reduction in the number of *foxi3a*-, *trpv6*- and *gcm2*-expressing cells in the neuromasts (Figure 5M and 5N). These data demonstrate that *foxi3a* plays a pivotal role in Nm ionocyte development. Interestingly, while *foxi3a* expression in skin ionocytes on the yolk was also nearly absent in *foxi3a* mutants (Supplementary Figure 4M), *gcm2*-expressing ionocytes are still present (Supplementary Figure 4N).

We next wondered whether the absence of Nm ionocytes in *foxi3a* mutants affects hair cell function. Since uptake of the vital dye FM1-43 depends on functional mechanotransduction (50–52), we tested whether hair cells in *foxi3a* mutants displayed decreased labeling with this dye. *foxi3a* mutants showed a slight reduction in FM1-43+ hair cells (Figure 5O) and hair cells displayed a significant reduction in FM1-43 intensity to 67% of average sibling’s levels (Figure 5P and 5Q). This demonstrates that mechanotransduction-dependent FM1-43 loading by hair cells is impaired in *foxi3a* mutants and that Nm ionocytes play an important role in regulating the ionic microenvironment surrounding lateral line hair cells.

## Discussion

The lateral line system is necessary for schooling behavior, mating, feeding and predator avoidance (Lush and Piotrowski, 2014). Therefore, it is crucial to maintain proper hair cell function, and regulate its microenvironment in the presence of ever-changing environmental factors (Hwang and Lee, 2007). We have made the unexpected discovery that pairs of skin-derived epithelial cells invade mature lateral line sensory organs post-embryonically. These cells are highly motile and, after invasion, rearrange and migrate extensively, before anchoring in stereotypical positions and differentiating into ionocytes.

Cell migration and tissue invasion are essential events driven by a diversity of signals and involved in many different processes such as development, regeneration, wound healing, immune system and lateral line formation itself (Kapsimali, 2017). However, these migratory events differ from the invasive cellular processes reported here in two major aspects: 1) they are restricted to developmental stages of organ formation; and 2) cells that contribute to mature organs during regeneration differentiate into sensory organ cells. For example, developing pulmonary neuroendocrine cells are sparsely distributed throughout the lung epithelium and then start to migrate leading to cell sorting and cluster formation (Kuo and Krasnow, 2015, Steinberg and Takeichi, 1994). Recruitment of cells into mature sensory organs occurs during taste bud regeneration when basal cells from the lingual epithelium surrounding the sensory organs regenerate taste cells (Barlow and Klein, 2015). In contrast, Nm ionocytes invade mature sensory organs while maintaining a distinct transcriptome.

### Signals that trigger Adaptive Cell Invasion and determine the Nm ionocyte lineage

The signals that attract epithelial cells to the sensory organs remain poorly understood. Taste bud cells secrete Shh which induces a response in the surrounding basal cells (Barlow and Klein, 2015). In case of Nm ionocytes, their invasion of neuromasts is not triggered by hair cell death and regeneration signals, instead they robustly respond to environmental changes, such as salinity and pH (Fig. 5E-J). Environmental changes also control ionocyte numbers in the skin via hormones such as cortisol, isotocin and prolactin (Chou et al., 2011, Evans et al., 2005, Flik and Perry, 1989, Guh et al., 2015, Kumai et al., 2015). The scRNA-seq analysis revealed Nm ionocytes indeed express the cortisol receptor nr3c1 (glucocorticoid receptor, GR), the main receptor involved in cortisol-induced ionocyte precursor differentiation, as well as the prolactin receptor prlra (Fig. 1Q) (Cruz et al., 2013a, Cruz et al., 2013b). Whether hormones trigger ACI needs to be further tested.

The identity of the signals that specify Nm ionocyte precursors among basal keratinocytes is unknown. Basal keratinocytes surrounding the neuromasts appear to be a heterogeneous population, as not all invade neuromasts upon salinity changes. In Medaka, neuromast border cells (nBCs) have been described to be induced by neuromasts (Seleit et al., 2017), however they are unlikely to be Nm ionocyte precursors as they do not enter the neuromasts. Nevertheless, it is likely that neuromasts induce ionocytes in their immediate vicinity. The transcription factors foxi3a/3b are required for skin ionocyte specification (Hsiao et al., 2007, Jänicke et al., 2007) and we find individual foxi3a+ cells in contact with neuromasts suggesting that these cells could be pre-invasion Nm ionocytes progenitors (Fig. S1T). foxi3a+ ionocyte progenitors in the developing skin are specified from epidermal stem cells by Notch-mediated lateral inhibition (Hsiao et al., 2007, Jänicke et al., 2007). Nm ionocytes also upregulate Notch signaling as they invade neuromasts. However, contrary to skin ionocyte specification, pharmacological inhibition of Notch signaling does not shift specification towards the ionocyte fate but abolishes pre-existing Nm ionocytes and prevents invasion of new cells. Future studies will shed light on the divergent roles of Notch signaling in early skin ionocyte specification and Nm ionocyte invasion.

### Tissue-specific ionocyte function

Ionocytes play crucial roles in the function of many organs. A rare ionocyte population was recently discovered in the lungs of mouse and humans and their loss is associated with cystic fibrosis (Montoro et al., 2018, Plasschaert et al., 2018). Different types of ionocytes in fish are functionally and molecularly analogous to ion transporting cells of the kidney of terrestrial vertebrates (Hwang and Chou, 2013). For example, HR ionocytes, specialized cells for acid/base regulation, share similarities with renal proximal tubular cells and type A-intercalated cells, all of which express carbonic anhydrases and H+-ATPases (Hwang and Chou, 2013, Skelton et al., 2010). Many kidney ion transporters are also highly expressed in the mammalian inner ear where they regulate the ionic composition of the endolymph (Lang et al., 2007). Consequently, mutations in a H+-ATPase subunit are not only associated with distal renal tubular acidosis, but also with progressive sensorineural hearing loss in humans and mouse models (Karet et al., 1999, Lorente-Canovas et al., 2013, Norgett et al., 2012). Similarly, mutations in Pendrin, a HCO3−/Cl− transporter, are associated with hearing loss caused by an acidification of the endolymph (Everett et al., 1997, Nakaya et al., 2007, Wangemann et al., 2007).

In the teleost ear ionocytes regulate the ionic composition of the endolymph (Becerra and Anadon, 1993, Mayer-Gostan et al., 1997). Even though the term ‘ionocyte’ has not been used to describe ion-regulating cells in the inner ear of non-teleost vertebrates, they possess functionally analogous cells. Possible homologs include K+ transporting dark cells in the vestibular system and marginal cells of the stria vascularis in the cochlea (Ciuman, 2009, Köppl et al., 2018). Vestibular dark cells regulate the endolymph in the ampullae of the semicircular canals and are adjacent to light cells, which were proposed to be homologous to ionocyte associate cells (Becerra and Anadon, 1993, Kimura, 1969, Köppl et al., 2018). In contrast to dark cells, light cells lack apical microvilli and send out membrane interdigitations to neighboring cells (Harada et al., 1989, Köppl et al., 2018). While microvilli-containing Nm ionocytes and their accessory cells appear morphologically similar to vestibular dark and light cells, respectively, detailed molecular and functional comparison between these cell types will be necessary to test homology with Nm ionocytes.

### Nm ionocyte function

Measurements of the endocupular potential in Xenopus laevis have suggested that the ionic composition of the cupula surrounding the hair cells is actively maintained (McGlone et al., 1979, Russell and Sellick, 1976). As Nm ionocytes are in direct contact with the cupula, we hypothesize that Nm ionocytes are crucial for this process. Lateral line hair cell function is impaired in acidified water and mutations that affect the development (merovingian) or function (persephone) of HR ionocytes result in mechanotransduction defects (Hailey et al., 2012, Lin et al., 2019, Stawicki et al., 2014). How HR ionocytes modulate hair cell function is not clear. It was proposed that loss of HR skin ionocytes in merovingian/gcm2 mutants leads to a global acidification of the extracellular environment that reduces hair cell mechanotransduction by affecting the endocupular potential or by impairing Ca2+ homeostasis (Horng et al., 2007, Horng et al., 2009, Stawicki et al., 2014). Our discovery of the previously undescribed Nm ionocytes that express multiple HR ionocyte genes and respond to acidic environmental conditions provides evidence that they play a role in acid/base regulation in the lateral line. Indeed, the loss of Nm ionocytes in foxi3a mutants leads to a decrease in mechanotransduction-dependent uptake of FM1-43 (Fig. 5P and 5Q). We propose that Nm ionocytes, which are located immediately adjacent to hair cells and are in direct contact with the hair cell microenvironment, are crucial for hair cell function.

### Nm ionocyte frequency is regulated by environmental changes and grants plasticity to the system

We have demonstrated that Nm ionocytes are recruited into neuromasts in a salinity- and pH-dependent manner, and we propose this is an adaptive trait that allows for tight control of hair cell microenvironment. This mechanism ensures sensory hair cell function and survival and subsequent acclimation in a wider range of environmental conditions. Acclimation is the process of facultative phenotypic changes in response to environmental cues. These changes can be reversible and repeatable during the organism’s lifetime and can affect different processes (Beaman et al., 2016). For example, tissue cell number has been observed to be environmentally regulated in the intestine of pythons during cycles of feeding and fasting or in murine adult stem cell compartments during long-term dietary interventions, low temperatures or light/dark cycles (Andrew et al., 2017, Casanova-Acebes et al., 2013, Ma et al., 2020, Mendez-Ferrer et al., 2008, Perry et al., 2019, Shwartz et al., 2020). Acclimation in the case of Nm ionocytes overcomes the need to make genetic changes that control the number of these cells per neuromast and confer higher plasticity within a generation. The plasticity of ionocyte numbers is not restricted to Nm ionocytes and has been observed for most ionocyte subtypes (Sardet et al., 1979, Varsamos et al., 2005). In addition, the ability of freshwater fish to respond to osmotic stress is also correlated with the capacity to modulate their ionocyte composition (Varsamos et al., 2005). It would be interesting to investigate whether fish that can inhabit larger salinity ranges (euryhaline fish) have an even more pronounced plasticity in their Nm ionocytes frequency compared to stenohaline fish, like zebrafish, that can only adapt to a smaller salinity range. ACI allows animals to cope with far from optimal environmental conditions. Even though this particular adaptation occurs within an individual in response to a narrow range of stimuli, it increases the range of environments in which the animal possesses high fitness. This extension of the geographical distribution that individual fish can inhabit due to ACI by ionocytes provides them with a selective advantage and is ultimately a driver of evolution.

## Conclusions

Even though lateral line organs have been studied since the 19th century, Nm ionocytes have eluded discovery. These newly described cells reported here are skin-derived and capable of invading mature sensory organs in response to environmental stimuli. *In vivo* time lapse imaging allowed us to characterize the migration of Nm ionocyte precursor cells from the skin into the lateral line. These studies also allowed us to observe and quantify the morphological changes these cells display during this highly dynamic process with high resolution, providing a detailed outline of the development of these newly discovered cells. Altogether, the comprehensive molecular, ultrastructural and functional characterization of Nm ionocytes provides evidence for their involvement in maintaining an ionic microenvironment that ensures robust hair cell mechanotransduction. Therefore, the discovery of ACI and Nm ionocytes not only underscores important roles for post-embryonic migration events in regulating the function of vertebrate mechanosensory organs, but also indicates that similar mechanisms may be associated with pathologies afflicting the inner ear.

## ACKNOWLEDGEMENTS

We are grateful to Drs. A. Sánchez Alvarado, R. Krumlauf, K. Si, R. Köster and the Piotrowski lab members for discussions and Drs. A. Sánchez Alvarado, A. Schier and N. Denans for critical reading of the manuscript. We also thank Drs. A. Schier, T. Carney, K. Poss, N. Lawson and B. Slaughter for reagents and fish lines, the Stowers Institute Aquatics Facility for fish care, J. Blanck and K. Ferro for cell sorting assistance, S. Baek, Y. Tsai, M. Harwood, S. Nowotarski and A. Peak for experimental assistance, C. Chen, D. Diaz, H. Li and J. Unruh for help with analysis and M. Miller for scientific illustrations. This work was performed to fulfill, in part, requirements for J. Peloggia’s thesis research in the Graduate School of the Stowers Institute. This work was funded by the Stowers Institute for Medical Research and the Hearing Health Foundation. Original data used for the results reported in this paper may be accessed from the Stowers Original Data Repository at *http://www.stowers.org/research/publications/LIBPB-15432020.*

## Author Contributions

Conceptualization, JP, DM and TP; Methodology, JP, DM, MEL; Investigation, JP, DM, PMG, ARC and MEL; Writing – Original Draft, JP, DM and TP; Writing – Review and Editing, PMG, ARC, MEL and YAP; Funding Acquisition, TP; Resources, YAP and TP; Supervision, TP.

---

**Supplementary Figure 1:**
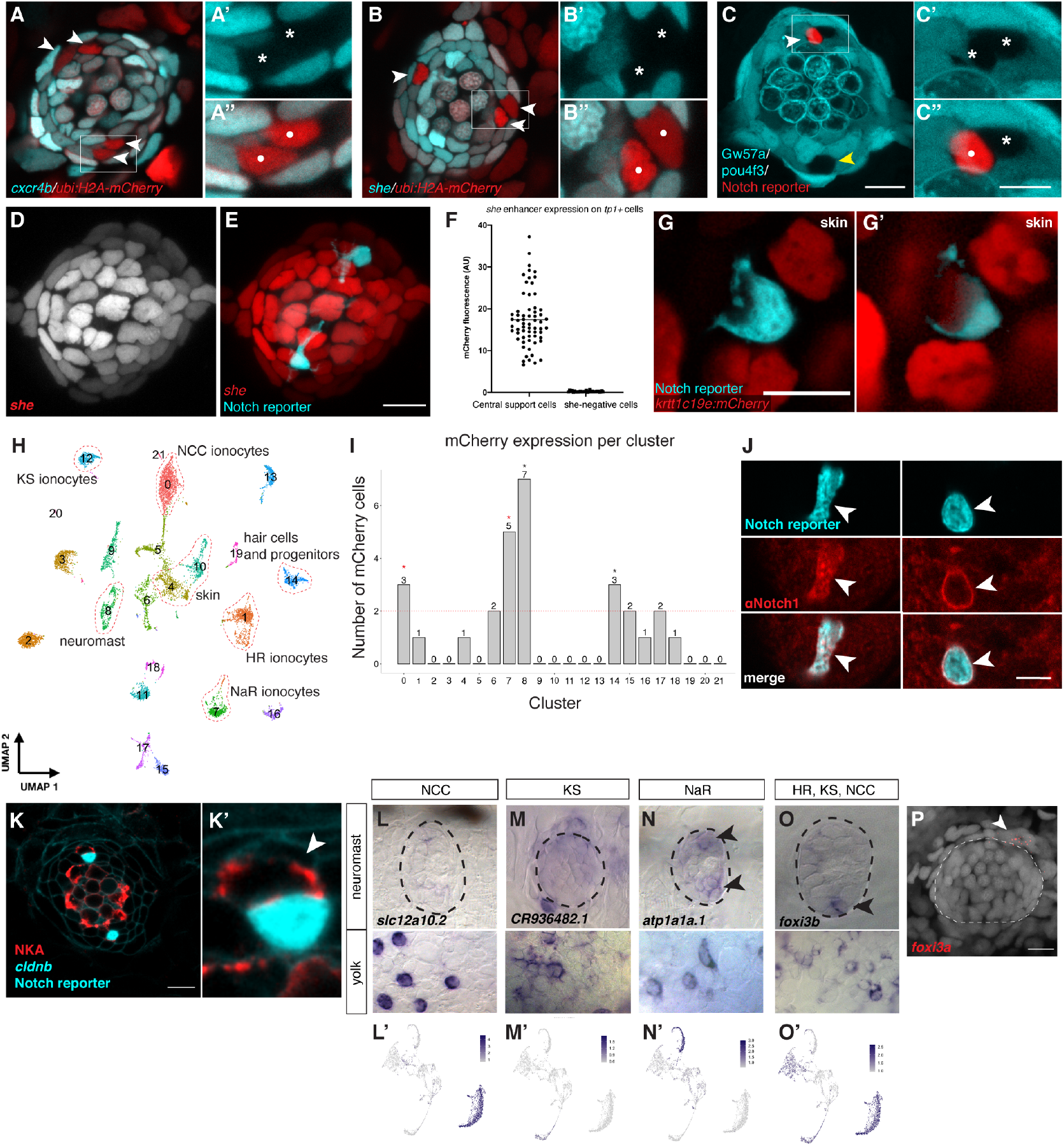
Non-lateral line cells do not express neuromast markers but share molecular features with ionocyte subpopulations. **(A), (A’) and (A”)** Neuromast of a 5 dpf cxcr4b-H2B-EGFP;ubi:H2A-mCherry larva containing two pairs of cells (A, arrowheads; A”, white dots) that are not labeled by the neuromast marker cxcr4b (A’, asterisks). Boxed area in (A) is shown at higher magnification in (A’) and (A”). **(B), (B’) and (B”)** Neuromast of a 4 dpf she:H2B-EGFP;ubi:H2A-mCherry larva containing pairs of cells (B, arrowheads; B”, white dots) that are not labeled by the neuromast transgenic ‘she’ line (B’, asterisks). Boxed area is shown at higher magnification in (B’) and (B”). **(C), (C’) and (C”)** Neuromast of a 5 dpf *Et(Gw57a:EGFP);Tg(pou4f3:GFP);tp1bglobin:mCherry* larva containing gaps in the fluorescence ((C), C’, asterisks). One gap contains a cell labeled by the Notch reporter ((C), white arrowhead; (C”), white dot), whereas the other gap does not contain a Notch+ cell due to mosaicism of the Notch reporter line ((C), yellow arrowhead). Boxed area is shown in higher magnification in (C’) and (C”). Scale bar of overview image = 10 μm, scale bar in magnified image = 5 μm. **(D)** Lateral line neuromast labeled by *she:H2A-mCherry.* **(E)** Merged image with *tp1bglobin:EGFP*. **(F)** Quantification of fluorescence intensity of all Notch+ cells in the neuromast shows that the newly discovered Notch+ cells in the poles do not express the H2A-mCherry driven by the lateral line specific enhancer she. **(G)** and **(G’)** Two individual z slices of the double transgenics *krtt1c19e:H2A-mCherry;tp1bglobin:EGFP* shows a rare pair of Notch+ and Notch− cell in the skin. **(H)** UMAP showing all clusters obtained by scRNA-seq after stand processing with Seurat v3.1 and quality control. Highlighted clusters represent different ionocyte populations, skin and lateral line cells. mCherry expression in clusters of **(H)**. We identified 35% mCherry cells in central support cells (clusters 8 and 14, black asterisks), and 29% of mCherry cells in ionocyte clusters (0 and 7, red asterisk). The remaining cells were distributed in low numbers in other clusters and were considered noise.**(J)** Notch+ (*tp1bglobin:EGFP*) non-lateral line cells express a Notch1 receptor, as shown by immunofluorescence with an anti-Notch1 antibody (arrowheads) Two individual z-slices of a confocal z-stack depicting the apical extension (left panel) as well as the cell body (right panel) are shown. Scale bar = 5 μm. (K, K’) Na+/K+ ATPase (NKA) presence in the Notch− cell and in hair cells. In situ hybridization of the NCC ionocyte cluster marker *slc12a10.2*. No staining is detectable in the neuromast poles (upper panel), but ionocytes on the yolk are labeled (lower panel). **(L’)** Feature plot of *slc12a10.2*. **(M)** In-situ hybridization of the KS ionocyte cluster marker *CR936482.1*. No strong staining is detectable in the neuromast poles (upper panel), but cells outside of the neuromast and ionocytes on the yolk are labeled (lower panel). **(M’)** Feature plot of *CR936482.1*. **(N)** In situ hybridization of the NaR ionocyte cluster marker *atp1a1a.1*. Weak staining is detectable in the neuromast poles (upper panel, arrowheads), and in presumed NaR ionocytes on the yolk (lower panel). **(N’)** Feature plot of *atp1a1a.1*. **(O)** In situ hybridization of the ionocyte marker *foxi3b* shows expression in the ventral pole of the neuromast. **(O’)** Feature plot of *foxi3b*. **(T)** HCR for *foxi3a* shows a Notch− *foxi3a* as well as a positive cell in the neuromast vicinity.

**Supplementary Figure 2:**
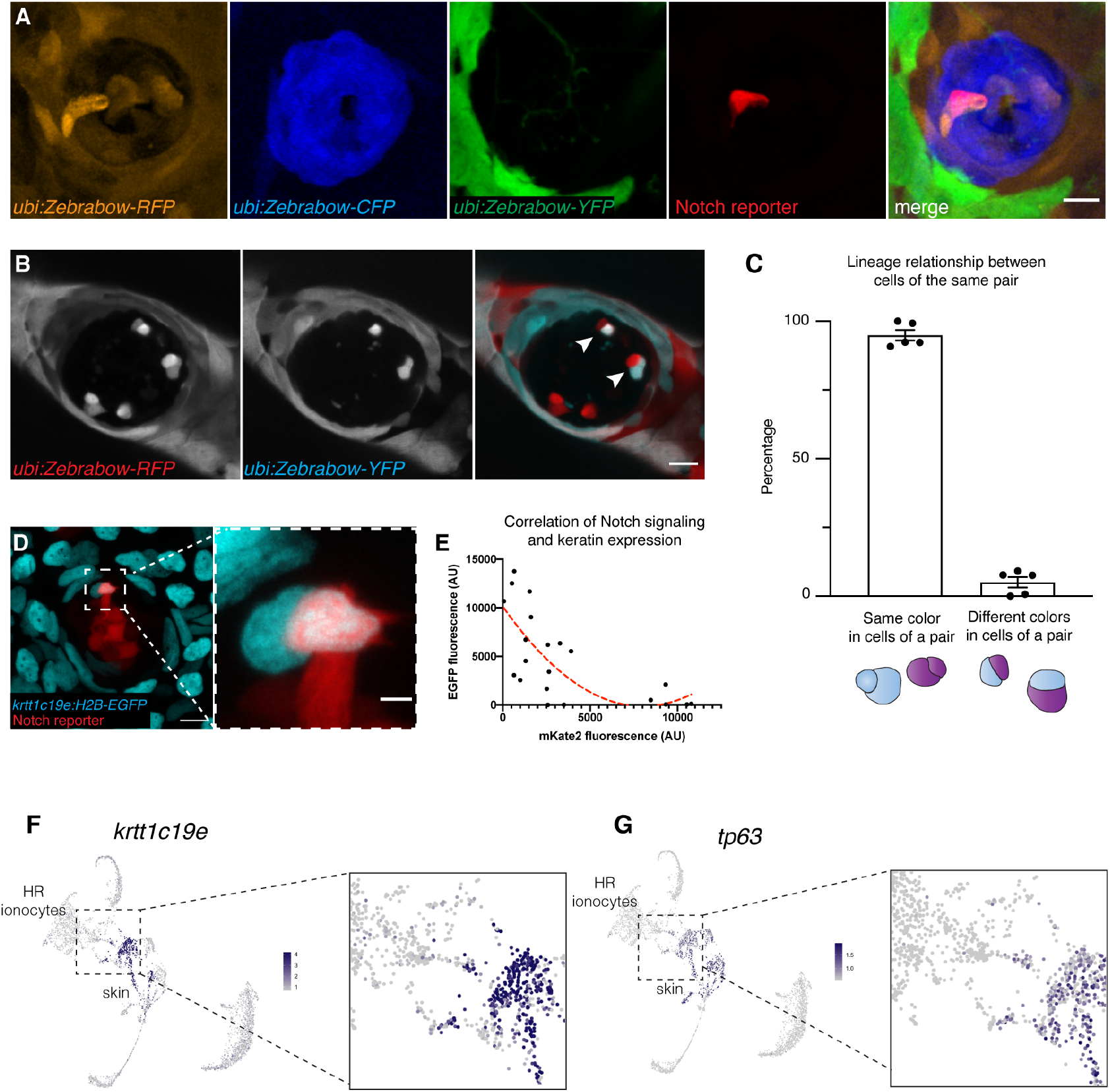
Nm ionocytes are derived from basal keratinocytes. **(A)** Analysis of a neuromasts in a 3-week old *ubi:Zebrabow;ubi:cre^ERt2^;tp1bglobin:mKate2* larva demonstrates that the cells with a different color hue than neuromast cells are the Notch+ cells. **(B)** Neuromast of 2-month old fish shows some pairs are composed of cells of different color hues, suggesting they did not originate from the same progenitor cell or the progenitor cells divided before Cre-induced recombination. **(C)** Ratio of ionocyte pairs that consist of cells of same or different color hues (n = 5 fish, 194 neuromasts). Error bars indicate SEM. **(D)** Neuromasts in a *tp1bglobin:mKate2;krtt1c19e:H2B-EGFP* larva show a pair of skin-derived cells in the neuromast, one of which is also labeled by the Notch reporter. **(E)** Inverse correlation (quadratic) between *krtt1c19e*-driven EGFP and *tp1bglobin*-driven mKate2 fluorescence suggests skin genes, such as *krtt1c19e* are turned off as Notch signaling increases. Adjusted R-squared: 0.4416 (n = 10 fish). **(F-G)** Feature plots for *krtt1c19e* (F) and *tp63* (G) show a possible differentiation trajectory from skin cells to HR ionocytes.

**Supplementary Figure 3:**
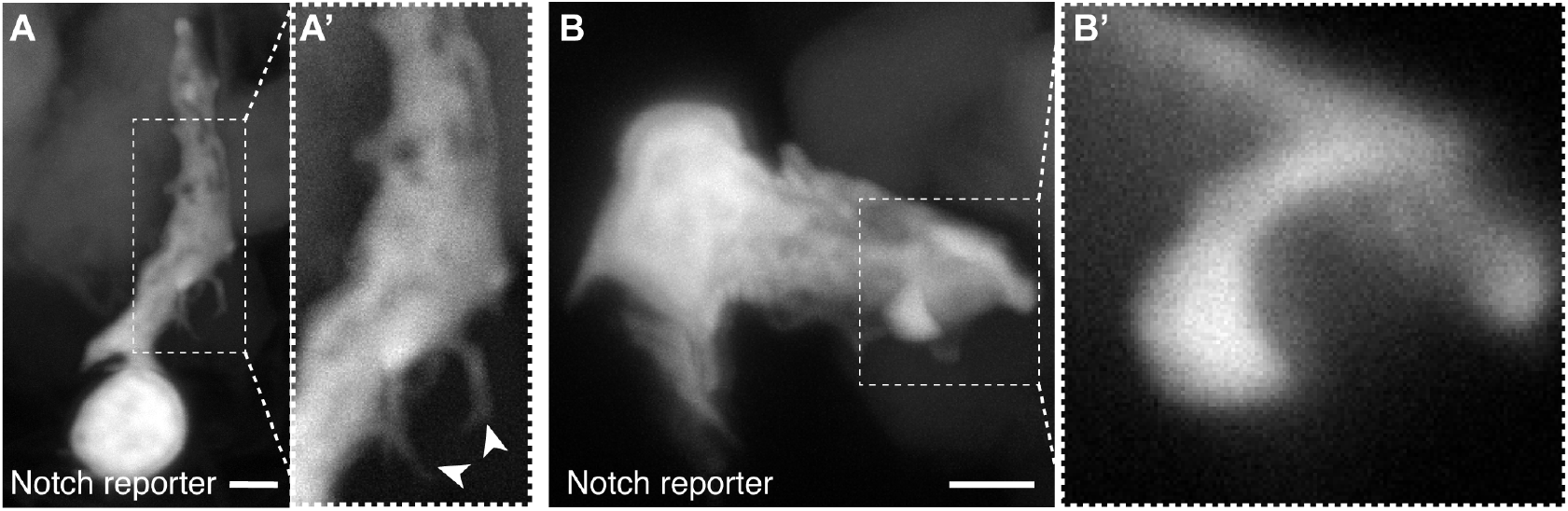
Nm ionocytes display multiple thin cellular protrusions and a pronounced apical extension. **(A-B’)** High-resolution confocal maximum projections of the apical extension formed by Notch+ cells (*tp1bglobin:EGFP*). (A) shows multiple thin cellular protrusions. **(A’)** Magnification of boxed area in (A) with white arrowheads pointing at protrusions. **(B)** depicts the most apical part of the extension of the Notch+ cell. **(B’)** magnification of the boxed area in (B) Scale bars = 2 μm. Images were gamma-adjusted to allow better visualization of fine cellular projections.

**Supplementary Figure 4:**
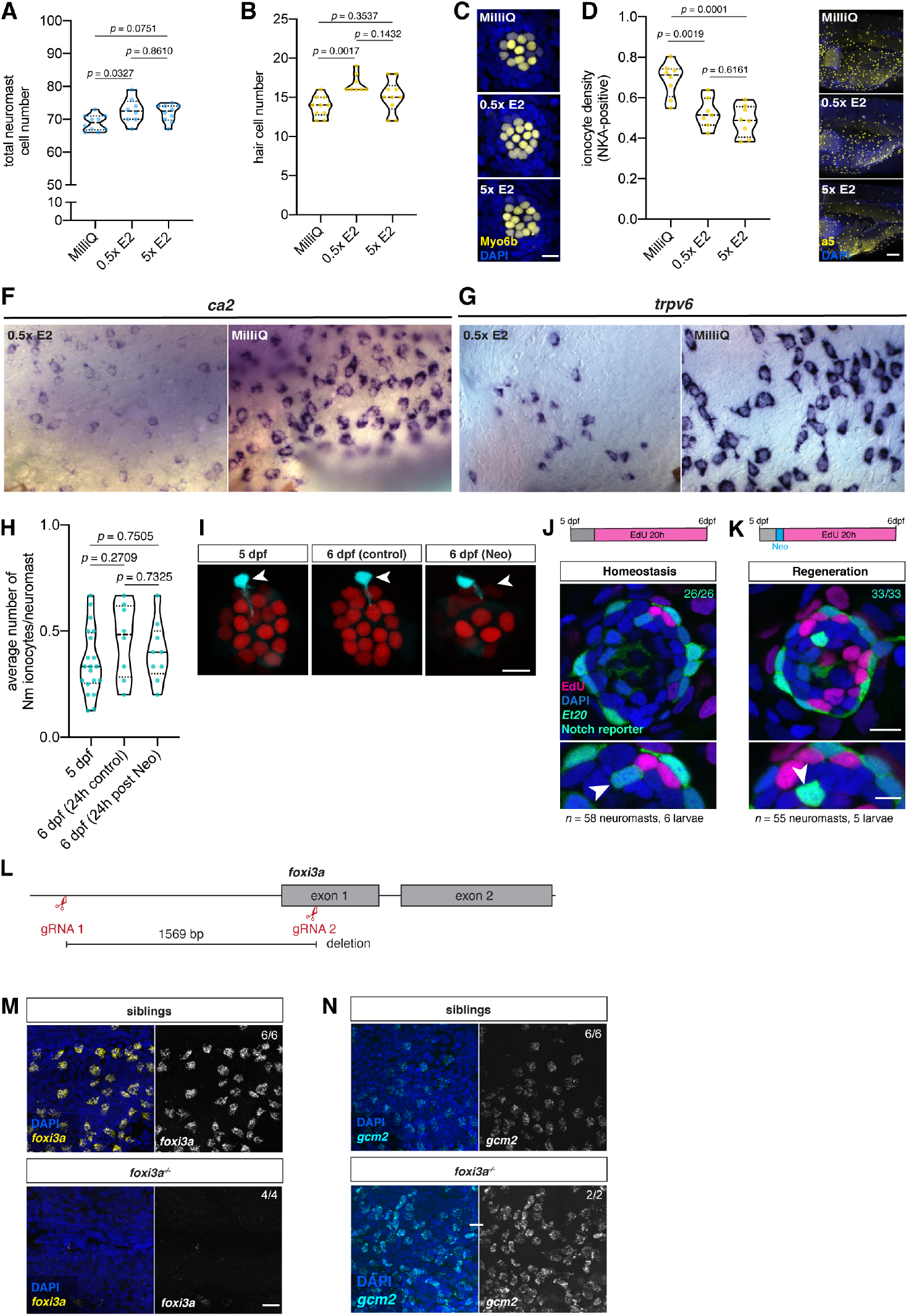
Salinity changes lead to tissue-specific phenotypes. **(A)** Incubation of larvae in MilliQ water (*n* = 5 fish, 10 neuromasts), but not 5x E2 (*n* = 5 fish, 10 neuromasts) from 3 dpf to 5 dpf significantly decreases the total neuromast cell number based on DAPI staining, compared to 0.5 x E2 media (*n* = 4 fish, 8 neuromasts; One-Way ANOVA with Tukey’s post hoc test). **(B)** Incubation of larvae in MilliQ water but not 5x E2 from 3 dpf to 5 dpf significantly decreases hair cell numbers compared to 0.5 x E2 media (*n* = 4 fish, 8 neuromasts; Kruskal-Wallis ANOVA with Dunn’s post hoc test). **(C)** Representative images of hair cell numbers in *Myo6b:H2B-Scarlet-I* larvae after incubation in media with different salinities, quantified in (B). Scale bar = 10 μm. **(D)** The density of ionocytes on the yolk detected with an anti-Na+-K+-ATPase (NKA) antibody increases after incubation in MilliQ water (*n* = 8 fish) but does not decrease significantly in 0.5x E2 (*n* = 7 fish) compared to 5x E2 media (*n* = 8 fish; One-Way ANOVA with Tukey’s post hoc test). **(E)** Representative images of NKA+ ionocytes on the yolk in conditions quantified in (D). Scale bar = 100 μm. (F) In situ hybridization depicting increased cell numbers of *ca2*-expressing ionocytes on the yolk after incubation of the larvae in MilliQ water for 48 h (right panel) compared to controls (left panel). **(G)** In situ hybridization depicting increased cell numbers of *trpv6* -expressing ionocytes on the yolk after incubation of the larvae in MilliQ water for 48 h (right panel) compared to controls (left panel). **(H)** Average Nm ionocyte numbers per fish do not significantly increase within 24 h after killing hair cells with the antibiotic neomycin (*n* = 20 fish, 305 neuromasts at 5 dpf; *n* = 8 fish, 119 control neuromasts at 6 dpf; *n* = 9 fish, 137 6 dpf neuromasts 24 h after neomycin treatment; One-Way ANOVA with Tukey’s post hoc test). **(I)** Representative images of neuromasts of *tp1bglobin:EGFP;Myo6b:H2B-Scarlet-I* larvae at 5 dpf before neomycin treatment (top panel), as well as 6 dpf control larvae (middle panel) and 6 dpf larvae 24 h post neomycin (lower panel). Scale bar = 10 μm. **(J)** EdU incorporation analysis of controls (*n* = 6 fish, 58 neuromasts with 26 Notch+ Nm ionocytes) and **(K)** Neomycin-treated neuromasts (*n* = 5 fish, 55 neuromasts with 33 Notch+ Nm ionocytes) of *tp1bglobin:EGFP;Et20* larvae counterstained with DAPI. Notch+ cells of Nm ionocytes do not divide during the first 20 h after Neomycin. **(L)** Schematic of the *foxi3a* promoter deletion strategy using CRISPR/Cas12a.**(M)** Expression of *foxi3a* on the yolk of *foxi3a*^−/−^ mutants (lower panel, n = 4 fish) and their siblings (upper panel, n = 6 fish) as detected by HCR. Scale bar = 20 μm. **(N)** Expression of *gcm2* on the yolk of *foxi3a*^−/−^ mutants (lower panel, *n* = 3 fish) and their siblings (upper panel, *n* = 12 fish) as detected by HCR. Scale bar = 20 μm.

## Supplementary Note 1: Methods

### Zebrafish lines and husbandry

All experiments were performed following the guidelines from the Stowers Institute IACUC review board. The published transgenic lines used were: *Tg(pou4f3:GAP-GFP)s356t* (89), *Tg(−8.0cldnb:lynEGFP)^zf106Tg^* (90), *Tg(tp1bglobin:EGFP)^um13^*, *Tg(tp1bglobin:hmgb1-mCherry)^jh11^* (33), *ET(krt4:EGFP)SqGw57a* (Kondrychyn et al., 2011), *Tg(ubi:creERt2)^cz1702Tg^* (91), *Tg(ubi:Zebrabow)^a131Tg^* (45), *TgBAC(cxcr4b:cxcr4b-GFP)^zf514Tg^* (92), *Tg(krtt1c19e:lyn-Tomato)^sq16Tg^* (31), *Tg(−8.0cldnb:H2A-mCherry)^psi4Tg^* (25), *Tg(krtt1c19e:H2A-mCherry)*. For generation of new transgenic lines, fusions of gene and fluorescent protein of interest were cloned into middle entry vectors (pME-MCS) using Gibson assembly (93) and recombined with targeted enhancers using the tol2 transgenesis kit (94). The following transgenic lines were made *Tg(ubi:H2A-mCherry)^psi18Tg^, Tg(she:H2B-EGFP)^psi59Tg^, Tg(krtt1c19e:H2B-EGFP)^psi63Tg^, Tg(she:H2A-mCherry)^psi57Tg^, Tg(tp1bglobin:lox-mKate2-lox-SpvB)^psi64Tg^, Tg(krtt1c19e:cre-MYC)^psi65Tg^, Tg(Myo6:H2B-mScarlet-I)^psi66Tg^* and *Tg(Myo6b:Lck-mScarlet-I)^psi67Tg^*.

### Generation of *foxi3a* mutants

CRISPR/Cas12a-mediated gene editing was performed to delete the *foxi3a* promoter and a part of the first exon (49). Two guide RNAs (gRNAs) targeting sequences 1377 bp upstream and 143 bp downstream of the start codon, respectively, were designed using DeepCpf1 (95) and checked for specificity and off-target effects by zebrafish genome BLAST (96) and CRISPRscan (49): gRNA 1: 5’-ATATGACAACCTACCGCAGA-3’; gRNA2: 5’-GGAGAATACGCAGGGCAGAC-3’. Tg(tp1bglobin:EGFP)um13 zebrafish one cell stage embryos were injected with 22.2 μM EnGen® Lba Cas12a (Cpf1) (New England Biolabs) and both gRNAs at a concentration of 13.8 mM each . All GFP+ larvae were raised to adulthood and germline-transmitting founders harboring the promoter deletion were identified by genotyping PCR with the following three primers: foxi3a-F: 5- CGATCAGAAAACGCCTGCAGACTGA-3’, foxi3a-R: 5’-GGGAGGTCTCACGAGTTTCATCAGATC-3’, foxi3a-R2: 5’- GCAACGAATGGAATCAGAATGTACAGTGC-3’. One F0 founder was outcrossed to wildtype fish to generate the F1 generation. F1 fish were raised to adulthood and heterozygous carriers of the deletion were identified by genotyping PCR. Multiple pairs of heterozygous F1 fish were incrossed and F2 larvae were analyzed for all experiments.

### In situ hybridization and Hybridization Chain Reaction

Whole mount in situ hybridization was performed according to http://research.stowers.org/piotrowskilab/in-situ-protocol.html. Probes were cloned and synthesized with the following primer pairs: foxi3b-F: TGA-CAATGGCAACTTCAGACGC, foxi3b-R: GCCCACAGAACTCAAAGGAG, slc12a10.2-F: CTGAAGCCAAACACACTTGTTTTGG, slc12a10.2-R: CTACATCTGCAGTAATACAATCATTGC, CR936482.1-F: AAACTTCAACTCTTCAAGAC, CR936482.1-R: ATCAAACGAGCTCCTTTATTC, trpv6-F: GGATTTTCTACATGACTCAGGAACC, trpv6-R: CAGTGATGGTGCACACCCTATATATTTG, ca2-F: CAGTTCTCTGGACTACTGGACATACC, ca2-R: GCATGCCTTCTATAGAATGTACGTTCAG. Reverse probes had a T3 polymerase sequence attached 5’ to allow for in vitro synthesis. PCR reaction was conducted with Phusion High Fidelity DNA polymerase (NEB, USA) and the product was purified by gel extraction according to manufacturer’s protocol (Monarch® DNA Gel Extraction Kit). In vitro transcription was carried out with T3 RNA Polymerase and DIG-labeling kit (Roche) for 2h at 37C, followed by 30 min of DNaseI treatment (Roche). RNA was purified with a LiCl Precipitation Solution (7.5 M) (AM9480, Invitrogen) and stored at −80C until use. Hybridization chain reaction (HCR) was performed according to manufacturer’s instructions for notch1b-B2, trpv6-B2, gcm2-B4, foxi3a-B1, krtt1c19e-B1 (38) (Molecular Instruments). The amplifiers used were B1-647, B2-546 and B4-488 (Molecular Instruments). HCR fish were subsequently stained with DAPI (5ug/mL) for 30 min at room temperature (RT) in the dark and washed three times with 5x SSCT before imaging.

### Immunohistochemistry

Larvae were anesthetized and fixed with 4% PFA overnight at 4C. Standard antibody staining was performed. Antibodies used were: TP63 (1:100, Santa Cruz, sc-8431), Anti-Notch1 intracellular domain rabbit polyclonal (1:100, Abcam, AB-306525), Monoclonal a5 ATPase, (Na (+) K(+)) alpha subunit (5 ug/mL, Developmental Studies Hybridoma Bank, AB-2166869), and rabbit anti-GFP (1:500, Invitrogen, A11122). Antibody staining was counterstained with DAPI (5ug/mL) for 30 min at RT in the dark and washed three times with PBST before imaging.

### Time-lapse and confocal imaging

Images were acquired using the confocal microscopes Zeiss LSM700, Zeiss LSM780 or Nikon Ti Eclipse with Yokogawa CSU-W1 spinning disk head equipped with a Hamamatsu Flash 4.0 sCMOS. Objective lenses used were Zeiss Plan-Apochromat 10x/0.45 M27 (air), Plan-Apochromat 20x/0.8 M27 (air) and LD C-Apochromat 40x/1.1 W Korr M27 (water) for the Zeiss Microscopes and a Nikon Plan Apo 40x 1.15NA LWD (water). Temperature was kept constant at 28.5C using Zeiss 780 standard incubation or a Stage Top Chamber (OkoLab) for the Nikon Microscope. For live imaging experiments, larvae were immobilized with tricaine (MS-222) up to 150 mg/L and mounted in glass bottom dishes (MatTek) with 0.8% low melting point agarose dissolved in 0.5x E2 with tricaine (100 mg/L). Laser lines used on the Zeiss confocal were Diode 405-30, Argon multiline laser (458, 488 and 514 nm), DPSS 561-10, HeNe 594 and 633 nm. A Nikon LUNV solid state laser launch was used for lasers 445, 515, and 561nm for CFP, YFP, and RFP respectively. Emission filters used on the Nikon were 480/30, 535/30, 605/70. All image acquisition was performed using Zen 2012 SP5 Black (Zeiss) and Nikon Elements AR 4.6 (Nikon) software.

### Electron Microscopy

#### Scanning Electron Microscopy

Zebrafish larvae (5 dpf) raised in 0.5x E2 were anesthetized and Nm ionocytes were mapped using a Zeiss LSM780 with LD C-Apochromat 40x/1.1 W Korr M27 (water) objective. Individually mapped larvae were fixed in 2.5% glutaraldehyde and 2% paraformaldehyde in PBS for 1 hour at room temperature on a rocker and stored at 4C until processing. Secondary fixation and staining with tannic acid, osmium, thiocarbohydrazide, and osmium were carried out as in (97). Samples were dehydrated in a graded ethanol series, critical point dried in a Tousimis Samdri 795 critical pint dryer, mounted on stubs, and imaged in a Hitachi TM4000 table top SEM at 10 kV.

#### Serial Block Face Scanning Electron Microscopy

Individual 5 dpf zebrafish larvae were analyzed for the presence of Nm ionocytes using the Notch reporter and then fixed in 2.5% glutaraldehyde, 2% paraformaldehyde, 1mM CaCl2, and 1% sucrose in 50 mM sodium cacodylate buffer for one hour at room temperature on a rocker, and processed for serial block face imaging in one of two ways: 1. (Figure 4B) As in (98), except that Hard Plus resin (Electron Microscopy Sciences) was used in place of Durcupan, or 2. (Figure 4C to 4F) With en bloc staining steps per (99–101), modified as follows and processed on the ASP-1000 automated sample processing robot (Microscopy Innovations): Buffered reduced osmium with formamide incubation was performed at room temperature for 4 hours, TCH incubation at 60 C for 45 minutes, incubation in 1% UA at room temperature for 4 hours plus an additional 2 hours at 60C, and lead acetate incubation for 2 hours at 60C. Samples were dehydrated in a graded series of acetone with 3- 100% exchanges for 15 minutes each step, and infiltrated with Hard Plus resin at 25% 50% and 75% for 30 minutes each and 3- 100% exchanges for 2 hours each. All samples were polymerized in either flat embed molds (Electron Microscopy Sciences) or using the minimal resin embedding technique as in (102). Once polymerized, samples were mounted on aluminum pins with silver conductive epoxy (Ted Pella, Inc). The minimal resin embedded sample was imaged in the Hitachi TM4000 SEM to verify the neuromast of interest for correlation with fluorescent imaging. Neuromast image volumes were acquired on a Zeiss Merlin with Gatan 3View 2XP at either or 2 kV, with slice thickness either 50nm or 80 nm, 10 nm pixel size and image acquisition area between of 55 μm or 65 μm. Charging was mitigated using focal charge compensation (Zeiss) with nitrogen set between 20% and 45%.

#### Tracing and Modeling

Tracing and modeling were performed in IMOD 4.9 (103). Final models were exported as wavefront objects and imported into Blender 2.83.4. Shade smooth was added to the objects and rendering was performed using the Blender Cycles Render at 29.97 frames per second. Intel AI Denoiser built-in on Blender was used for animation post-processing and denoising.

### Single Cell RNA sequencing

#### Embryo dissociation and FACS

A total of 600 larvae 5 dpf Tg(−8.0cldnb:lynEGFP);Tg(tp1bglobin:hmgb1-mCherry)jh11 were screened for double positive fluorescence. 300 embryos were dissociated in 4.5 mL of 0.25% trypsin-EDTA (Thermo Fisher Scientific, Waltham, MA. USA) by pipetting them for 2.5 min on ice with 1uM Actinomycin D (Sigma A1410) to inhibit new transcription. Cells were stained with DAPI (5 μg/mL) and Draq5 (1:1000) (Thermo) and sorted for DAPI-negative and Draq5-positive cells for viability control. Cells were sorted into 90% MeOH for fixation. Cell sorting was performed by the Cytometry Facility of the Stowers Institute for Medical Research using the BD Influx Cell Sorter (BD Biosciences, San Jose, CA. USA).

#### 10X Chromium scRNA-seq library construction

MeOH-fixed cells were rehydrated with rehydration buffer (0.5% BSA and 0.5 U/μl RNase-inhibitor in ice-cold DPBS) following manufacturer’s instructions (10X Genomics). Approximately 15.000 cells were loaded into the Chromium Single Cell Controller (10x Genomics). For library preparation, Chromium Next GEM Single Cell 3’ GEM, Library Gel Bead Kit v3.1 was used. The sample concentration was measured on a Bioanalyzer (Agilent) and sequenced with NextSeq 500 High Output Kit v3 with read length of 28 bp Read 1, 8 bp i7 index and 91 bp Read 2 (150 cycles) (Illumina).

#### Read Alignment and Quantification and Quality Control

Reads were de-multiplexed and aligned to version Ensembl GRCz11 of the zebrafish genome using the CellRanger (v3.0.0) pipeline. Following analysis of UMI count matrix was performed using Seurat (version 3.1.2) (34, 104). Initial quality control filtered out all genes expressed in less than three cells, less than 250 genes, more than 30% mitochondrial genes and 30.000 UMIs. Standard Seurat v3.1.2 workflow was performed, following the steps from the pbmc3k online tutorial (https://satijalab.org/seurat/v3.2/pbmc3k_tutorial.html). Dimensional reduction, resolution and the final UMAP were then validated by comparing their biomarkers, differentially expressed genes obtained with the function FindAllMarkers(), to known markers based on literature. Further cluster subset was performed for all detected ionocytes and skin cells (defined as tp63+). Analysis code is available at: https://github.com/Piotrowski-Lab/scRNAseq_Nm_ionocytes_analysis_Seuratv3_Peloggia_Muench_et_al_2020

#### Acclimation of zebrafish larvae to different pH and salinity environments

For the acclimation experiments, 0.1x, 0.5x, 2.5x, 5x E2 dilutions of E2 media were prepared with distilled water from a 20x E2 stock solution (17.5g/L NaCl, 0.75g KCl, 4.9g/L MgSO4•7H2O, 0.41g/L KH2PO4, 0.142g/L Na2HPO4, 2.9g/L CaCl2•2H2O, pH = 6.0-6.1). To prepare pH 4, 7, and 10 media, 0.5x E2 medium was supplemented with 300 μM MES (Chem-Impex), 300 μM MOPS (Sigma), or 300 μM tricine (Sigma), respectively (modified from (48)). In all experiments, larvae were first raised in regular 0.5x E2 medium to 3 dpf. Between 3 and 5 dpf, the larvae were then incubated in diluted E2 media or media with different pH for 48 h. Embryo medium was exchanged daily to ensure stable pH or salinity. No significant differences in larvae survival were observed between the different conditions. MilliQ water parameters are conductivity of μS/cm, salinity of 0.0 ppm, alkalinity of 17.5 ppm (as CaCO3) and general hardness of 7.5 ppm (as CaCO3) and a pH of 7.6. Conductivity and salinity were measured with YSI 30 (YSI incorporated), and alkalinity, general hardness and pH were measure with SpinTouch FF104 (LaMotte).

#### Quantification of Nm ionocyte frequency

To quantify Nm ionocyte frequency based on the absence *Et(Gw57a:EGFP)/Tg(pou4f3:GFP)* fluorescence, 5 dpf larvae were embedded in 0.8% low melting point agarose and gaps in *Et(Gw57a)* fluorescence were counted using a Zeiss LSM780 confocal. A gap in the fluorescence in the neuromast poles that was fully separated from other fluorescence gaps was considered to be one Nm ionocyte, regardless of the number of cells within the gap. A total number of 14-16 neuromasts of the anterior lateral line (Ml1, O2, Ml2, IO4, O1; for nomenclature see (105)), as well as the posterior lateral line (L1-L9, LII.1, LII.2; for nomenclature see (26)), were quantified per larva, unless otherwise noted. To quantify Nm ionocyte frequency based on the Notch-reporter (*tp1bglob:EGFP*) fluorescence, larvae were anesthetized with 150 mg/L tricaine (MS-222) and juvenile and adults were killed using cold-shock protocol. The fish were embedded in 0.8% low melting point agarose and Notch+ cells were counted on a confocal microscope. All cells in the neuromast polar compartments showing strong labeling by the Notch reporter and/or displaying pronounced apical protrusions were included in the analysis. A total number of 14-16 neuromasts of the anterior lateral line (Ml1, O2, Ml2, IO4, O1), as well as the posterior lateral line (L1-L9, LII.1, LII.2), were quantified per larva, unless otherwise noted.

#### Sensory hair cell ablation

To ablate mechanosensory hair cells, 5 dpf *tp1bglob:EGFP* larvae were incubated in 300 μM neomycin (Sigma-Aldrich) for 30 min (106). After the antibiotic was washed out, larvae were allowed to recover in 0.5x E2 for 24 h and Nm ionocyte number was quantified based on Notch reporter fluorescence as described above.

#### Proliferation analysis

5 dpf tp1bglob:EGFP;Et20 larvae were incubated in 3.3 mM EdU (Carbosynth) in 1% DMSO in E2 medium for 20h following Neomycin-induced sensory hair cell ablation (see above). EdU staining, anti-GFP immunohistochemistry and DAPI counterstaining were performed as previously described (Lush et al., 2019).

#### Manipulation of Notch signaling

Notch signaling was inhibited by incubating larvae in 50 μM of the gamma secretase inhibitor LY411575 (Selleckchem) with 0.25% DMSO in 0.5x E2 media for 24 h. Control larvae were incubated with 0.25% DMSO in 0.5x E2 media only. Nm ionocyte frequency was quantified based on gaps in *Et(Gw57a:EGFP)/Tg(pou4f3:GFP)* fluorescence as described above. Three neuromasts of both the anterior lateral line (Ml1, O2, Ml2) and the posterior lateral line (L1, L2, L3) were quantified per larva.

#### FM1-43 dye uptake

To analyze mechanotransduction-dependent uptake of FM1-43FX by hair cells, individual 5 dpf larvae were incubated with 2.25 μM FM1-43FX (Invitrogen, F35355, diluted in 0.5x E2 medium) for 1 min, washed two times with 0.5x E2 containing 200 mg/mL MS-222 for 15s and embedded in a small drop of 0.8% low-melting point agarose supplemented with 140 mg/mL on a glass bottom dish. After letting the agarose solidify for 1 min, the L2 and L3 neuromasts of each larva were imaged as a z-stack of 25 sections (1 μm Z step).

### Image Analysis

All image analyses were performed in Fiji (107) and 3D renderings in Imaris (Bitplane).

#### Nonlinear operations (gamma)

The *she* (lateral line) and *tp1bglobin* (Notch reporter) enhancers drive expression at different ranges in different cells. Therefore, non-linear changes (gamma) were applied to make all features visible. Gamma ranged from 0.5 to 0.8 and was applied whenever stated in figure legends. Image quantification was always performed with the raw file and never with gamma-modified images.

#### Position quantification

Cell positions for mantle cells and Notch+ Nm ionocytes were quantified in Fiji or Imaris software, and calculations were performed as previously described (25, 106).

#### Linear unmixing

For proper separations of *ubi:Zebrabow* and mKate2 fluorescent proteins, linear unmixing was performed. Fluorescent proteins were excited by using 458 laser line for CFP, 514 for YFP and RFP and 594 for mKate2. Z stacks were acquired on a Zeiss 780 confocal set on lambda mode and 8.9nm resolution. Images were unmixed using a collected reference spectrum for mKate2 and by manually detecting CFP, YFP and tdTomato spectra on Zen Black (Zeiss) based on FPBase reference spectra (108).

#### Tracking of individual cells of migrating Nm ionocyte cell pairs

To visualize the path of both cells of the migrating Nm ionocyte precursor pair over time, confocal time lapse recordings were analyzed using the mTrackJ plugin (109) in ImageJ and cell nuclei tracked through individual z-planes. Tracks were then visualized on maximum projections of the time lapse recording using the “Movie” function in mTrackJ. The distance to the start point of each track (D2S) was obtained using the “Measure” function in the mTrackJ plugin.

#### Quantification of migrating Nm ionocyte precursor speed

To quantify migrating Nm ionocyte precursor speed, z-stacks of confocal time lapse recordings were maximum projected and corrected for tissue drift using the StackRegJ plugin (110). The nucleus of the Notch+ cell was then tracked based on *cldnB:H2A-Cherry* or *ubi:H2A-Cherry* signal and using *tp1bglob:EGFP* fluorescence as an additional reference. The summed distance of the entire track, as well as the net distance between the first and the last tracking point were calculated using the “Measure” function of the mTrackJ plugin (final track length and final D2S, respectively) and the speed calculated by dividing the distance by the elapsed time.

#### Quantification of FM1-43 fluorescence intensity

To quantify FM1-43 dye uptake by hair cells, maximum intensity projections of 12 z-planes of the acquired z-stacks were generated, starting just below the cuticular plate and encompassing the entire cell body of the hair cells. A mask was then drawn around the labeled hair cells and the average fluorescence intensity calculated using the ‘Measure’ function. Four identical masks were drawn in multiple regions within close vicinity of the hair cells to calculate the average background intensity. Data is shown as hair cell fluorescence intensity/background, normalized to the average of the siblings and shown as percentage.

### Statistical Analysis

All statistical tests were performed using GraphPad Prism 8 (version 8.4.3) as indicated in the figure legends. Normal distribution was assessed using Kolmogorov-Smirnov test. When comparing data from more than two groups, statistical significance was calculated using either one-way ANOVA with Tukey’s post hoc test or non-parametric Kruskal-Wallis ANOVA with Dunn’s post hoc test. Data from two groups were compared using two-tailed unpaired t-test or two-tailed Mann–Whitney U test. Binomial analysis was performed in R (111, 112). p-values and sample size per experiment are individually specified for each experiment in the figure legends. Plots were made in Microsoft Excel (version 16.39), GraphPad Prism 8 and the R package ggplot2 (113).

